# Solution structure of NPSL2, a regulatory element in the oncomiR-1 RNA

**DOI:** 10.1101/2022.03.27.485969

**Authors:** Yaping Liu, Aldrex Munsayac, Ian Hall, Sarah C. Keane

## Abstract

The miR-17~92a polycistron, also known as oncomiR-1, is commonly overexpressed in multiple cancers and has several oncogenic properties. OncomiR-1 encodes six constituent microRNAs (miRs), each enzymatically processed with different efficiencies. However, the structural mechanism that regulates this differential processing remains unclear. Chemical probing of oncomiR-1 revealed that the Drosha cleavage sites of pri-miR-92a are sequestered in a four-way junction. NPSL2, an independent stem loop element, is positioned just upstream of pri-miR-92a and sequesters a crucial part of the sequence that constitutes the basal helix of pri-miR-92a. Disruption of the NPSL2 hairpin structure could promote the formation of a pri-miR-92a structure that is primed for processing by Drosha. Thus, NPSL2 is predicted to function as a structural switch, regulating pri-miR-92a processing. Here, we determined the solution structure of NPSL2 using solution NMR spectroscopy. This is the first high-solution structure of an oncomiR-1 element. NPSL2 adopts a hairpin structure with a large, but highly structured, apical and internal loops. The 10-bp apical loop contains a pH-sensitive A^+^·C mismatch. Additionally, several adenosines within the apical and internal loops have elevated p*K*_a_ values. The protonation of these adenosines can stabilize the NPSL2 structure through electrostatic interactions. Our study provides fundamental insights into the secondary and tertiary structure of an important RNA hairpin proposed to regulate miR biogenesis.

## Introduction

MicroRNAs (miRs) are involved in the post-transcriptional regulation of gene expression in a wide variety of cellular processes from diverse organisms.^1–6^ miRs are found within long primary miRNA (pri-miR) transcripts, which may contain one (monocistronic) or several (polycistronic) miR sequences. These pri-miRs are processed into ~70 nucleotide (nt) precursor miRs (pre-miRs) by Drosha and its cofactor DiGeorge syndrome critical region (DGCR8) in the nucleus.^7–9^ The pre-miR is exported from the nucleus to cytoplasm by exportin 5 (XPO5) and further processed by Dicer, into ~22 nt RNA duplex.^10–12^ One strand is then incorporated into the RNA-induced silencing complex (RISC) which targets a messenger RNA (mRNA) with a complementary sequence.^13–15^

OncomiRs are miR genes that function as oncogenes and their expression are tightly related with the development of myriad cancers.^16^ The human miR-17~92 cluster is strongly linked to oncogenesis^17^ and is also known as oncomiR-1. Expression of oncomiR-1 has significant effects on tumorigenesis and development in mouse model system.^18^ Abnormal expression of oncomiR-1 is involved in the process of many cancers, including lung, breast, colon, prostate, thyroid, pancreas, stomach, hepatocellular and lymphoma.^19–23^ *In vitro* studies have shown that both c-MYC and E2F are transcription factors that have effects on cancer.^24–26^ The expression of oncomiR-1 in a mouse model shows that this cluster is essential; the knockout of oncomiR-1 in newborn mice resulted in death soon after birth.^18^ These findings reflect the large impact of oncomiR-1 on gene regulation, development and disease.

OncomiR-1 is an 827 nt long transcript that is located at the *13q31.3* genomic locus.^27^ Its secondary structure has been probed using dimethyl sulfate (DMS) and selective 2’-hydroxyl analyzed by primer extension (SHAPE) chemistries (**Fig. S1**).^28^ The oncomiR-1 primary transcript is polycistronic, containing six different miRs (miR-17, miR-18a, miR-19a, miR-20a, miR-19b and miR-92a), which are co-transcribed. However, the relative abundance of the mature miRs processed from oncomiR-1 is altered under different physiological conditions.^29,30^ Aberrant processing of oncomiR-1 disrupts the balance of mature miRs produced, leading to pathological conditions such as cancers^18^ and developmental deformities.^31^ For example, one of the constituent miRs, miR-92a, promotes tissue growth and angiogenesis and its overexpression is related to erythroleukemia,^32^ colorectal cancer,^33^ and lung cancer.^33,34^ Conversely, miR-17 functions as a tumor suppressor in breast cancer by inhibiting amplified in breast cancer 1 (*AIB1*) mRNA translation.^35^ Furthermore, some miRs play dual roles. For example, miR-18a can function as an oncogene, promoting cancer progression;^36^ and also as a tumor suppressor in ovarian^37^ or lung cancer.^38^ The contrasting functions of the miRs within oncomiR-1 highlight the importance of precise regulation of the biogenesis of these miRs from the oncomiR-1 transcript.

Previous studies showed that oncomiR-1 adopts a compact and globular tertiary structure, which may provide an explanation for the differential processing of its constituent miRs.^39,40^ The 3’-core domain of oncomiR-1 (pre-miR-19b through pre-miR-92a) is internalized within the folded oncomiR-1 structure. The sequestration of these 3’-core elements results in Drosha cleavage sites that are buried and therefore processed less efficiently.^40,41^

NPSL2, a non-precursor miR element, is located within the 3’-core domain, positioned between pre-miR-19b and pre-miR-92a. Chemical and enzymatic probing of NPSL2 revealed that NPSL2 adopts a hairpin structure with purine-rich internal and apical loops, consistent with the secondary structure of full-length oncomiR-1.^28,41^ NPSL2 is reported to form tertiary contacts with pre-miR-19b that contributes to the compact folding of oncomiR-1 and repression of pri-miR-92a processing.^41^ Disruption of the tertiary contacts between NPSL2 and pre-miR-19b through mutagenesis of the unpaired adenosines in the internal loop of NPSL2 also enhanced pri-miR-92a processing.^41^

The secondary structure of oncomiR-1 (**Fig. S1**) reveals that pre-miR-92a adopts a hairpin structure lacking a basal helix, its Drosha cleavage sites sequestered in a four-way junction.^28^ To promote processing of pri-miR-92a, there may exist a conformational rearrangement that promotes formation of the basal helix through remodeling of the NPSL2 element. NPSL2 therefore, may serve as a linchpin for the conformational rearrangements that regulate pri-miR-92a processing through tertiary interactions or structure remodeling. Additional trans-acting factors, like RNA binding proteins, may contribute to the remodeling of the 3’-core domain, further regulating pri-miR-92a processing.

In this work, we determined the solution structure of NPSL2 using a combination of NMR spectroscopy and small angle X-ray scattering (SAXS). We found that the internal and apical loops of NPSL2 were highly-organized, with extensive base stacking interactions. This represents the first high-resolution structure of an oncomiR-1 element. We used circular dichroism spectroscopy to analyze the thermal stability of various NPSL2 constructs under different solution conditions. Through analysis of NMR data, we identified several adenosines within the apical and internal loops that have significantly shifted p*K*_a_ values. Protonation of these adenosines can stabilize the structure of NPSL2 through formation of an A^+^·C mismatch within the apical loop, phosphate-base, and cation-π interactions.

## Results

### NPSL2 adopts a hairpin structure

WT NPSL2, a 37 nt long domain within the oncomiR-1 RNA that is predicted to fold into a hairpin structure (**Fig. 1A**), is highly conserved among different species.^28^ To facilitate *in vitro* transcription and stabilize the hairpin structure, 3 G-C base pairs were added to the base of the hairpin (**Fig. 1B**). The addition of the three G-C base pairs to the WT NPSL2 sequence stabilized the RNA by ~13 °C in a thermal denaturation assay (**Fig. 1C**). Additionally, NMR analysis revealed that the overall structure of the RNA was not affected by the additional base pairs. ^1^H-^1^H NOESY and ^1^H-^13^C HSQC spectra of both NPSL2 and WT NPSL2 display similar structural features, with minor chemical shift perturbations localized to nucleotides in the base of the helix (**Fig. 1D**). Due to the similarities in structures and the increased stability of the hairpin, the G-C-containing NPSL2 sequence was used as the basis for further structural studies. Analysis of the G-C-containing NPSL2 by native gel and size-exclusion chromatography shows that the NPSL2 structure is homogenous in solution (**Fig. S2A,B**).

**Fig. 1.**
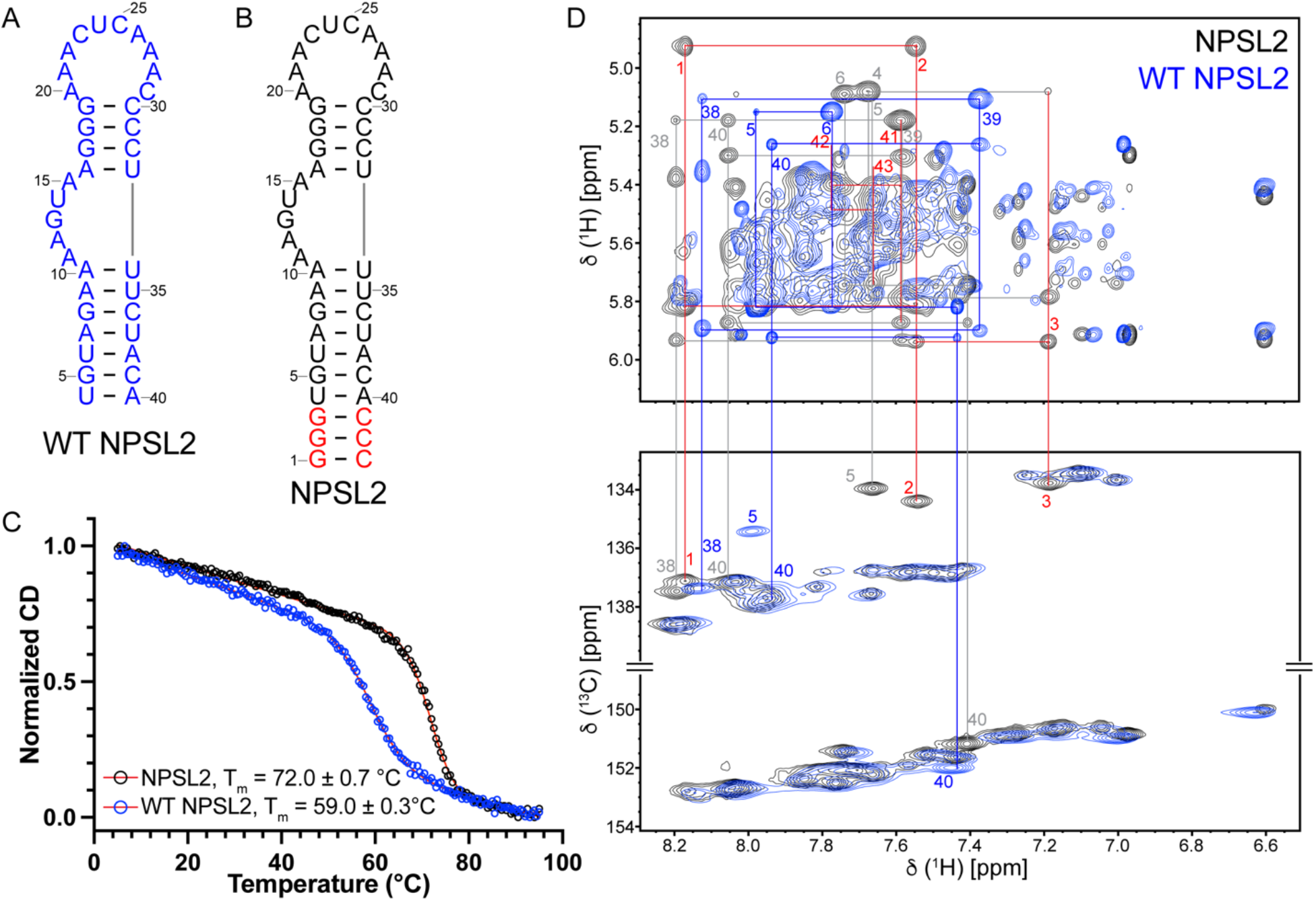
NPSL2 adopts a hairpin structure. (A) Sequence and secondary structure model of WT NPSL2. (B) Sequence and secondary structure model of NPSL2. (C) Normalized CD-thermal denaturation curves of WT NPSL2 (blue) and NPSL2 (black). (D) ^1^H-^1^H NOESY (top) and ^1^H-^13^C HSQC (bottom) spectra of WT NPSL2 (blue) and NPSL2 (black). Select chemical shift assignments are shown. Resonances that are unique to NPSL2 are denoted with red labels, with flanking sequences denoted in grey. Blue labels denote resonances from the first three base pairs of WT NPSL2. Secondary structures were rendered using RNA2Drawer.^88^

### Oligonucleotide-based approach to facilitate resonance assignments of NPSL2

The 2D ^1^H-^1^H NOESY spectrum of NPSL2 exhibited severe signal overlap and was unassignable using a full-length, unlabeled sample. To reduce the spectral complexity, we designed employed a “divide-and-conquer” approach, where we used smaller RNA oligonucleotides representing distinct structural elements of the full-length RNA (**Fig. S3**). We designed two oligonucleotides: NPSL2_Frag1 and NPSL2_Frag2 (**Fig. S3A**). NPSL2_Frag1 is comprised of the A-helical stem at the base of NPSL2 and is capped with a non-native GAGA tetraloop (**Fig. S3A**). The GAGA tetraloop is structurally stable and yields characteristic and unambiguous signals in the 2D ^1^H-^1^H NOESY spectra, which serve as an indicator of proper folding of the RNA and facilitate resonance assignments.^42,43^ The observed NOE signals of NPSL2_Frag1 revealed that it adopted an A-form helical stem with a properly folded GAGA tetraloop (**Fig. S4**).

NPSL2_Frag2 was designed to isolate the resonances from the apical loop of NPSL2. The NPSL2_Frag2 oligonucleotide consists of the apical stem and upper helical region. The construct was extended by two G-C base pairs at the base of the helix, to facilitate efficient transcription and to stabilize the helical structure (**Fig. S3A**). Six out of ten nucleotides in the apical loop are adenosines, which are largely degenerate in chemical shift. However, the small size of NPSL2_Frag2 enabled complete resonance assignments (**Fig. S5**).

The nonexchangeable signals for NPSL2_Frag1 and NPSL2_Frag2 were unambiguously assigned based on the analysis of 2D ^1^H-^1^H NOESY, 2D ^1^H-^1^H TOCSY, and 2D ^1^H-^13^C HMQC spectra (**Figs. S4, S5**). All assignments are in good agreement with predicted chemical shifts (**Figs. S4, S5**). Resonance assignments have been deposited in the BMRB (NPSL2_Frag1: 51348, NPSL2_Frag2: 51349). Notably, this study also contributed a number of previously uncharacterized chemical shifts to the RNA chemical shift database (see open circles in **Figs. S4, S5**).

Comparison of the ^1^H-^1^H NOESY spectra for NPSL2, NPSL2_Frag1, and NPSL2_Frag2 revealed that the oligonucleotide fragments adopt the same conformation in isolation as within the full length NPSL2 (**Fig. S3B**). This oligonucleotide-based approach provided the foundation for chemical shift assignments for the full-length NPSL2 RNA. With these data, we were able to unambiguously assign the resonances from the bottom and upper stems of NPSL2, however, the internal and apical loops assignments remained unresolvable.

### NMR assignments of NPSL2 by ^2^H-editing approach

Even with the assigned oligonucleotides as a guide, the stretches of adenosines in the apical and internal loops of NPSL2 were unassignable in the fully protonated ^1^H-^1^H NOESY spectrum due to the existence of broad, overlapping cross peaks. To reduce spectral complexity, we implemented a deuterium-edited approach, that involved the analysis of ^1^H-^1^H NOESY spectra collected on NPSL2 RNAs prepared with different combinations of fully protiated, fully deuterated, and partially deuterated nucleotide triphosphates. The following NPSL2 samples were prepared: A^2r^G^r^C^r^, A^2r^G^r^, A^8^C^H^, A^H^C^H^, A^H^G^8^, G^H^, A^2r^U^r^, A^8^U^6r^; superscripts denote sites of protiation on a given nucleoside, all other sites deuterated; e,g., G^H^ = fully protiated guanosines, A^2r^ = adenosines protiated at the C2 and ribose carbons.

We initially conducted ^1^H-^1^H NOESY experiments at 37 °C using an A^2r^G^r^-labeled NPSL2 RNA. This spectrum revealed only 5 adenosine H2 signals out of a total of 15 expected adenosine H2 signals (**Fig. S6A,B**). By reducing the temperature from 37 °C to 15 °C, most of the H2 signals reappeared in the ^1^H NMR spectrum (**Fig. S6C**). Therefore, all subsequent NMR data for structural analysis were collected at 15 °C.

^1^H-^1^H NOESY spectra of selectively deuterated NPSL2 RNAs exhibited good resolution and sensitivity (**Fig. 2A**). Sequential and long-range NOEs involving the adenosine-C2 protons were assigned primarily from the spectra obtained on A^2r^U^r^- and A^2r^G^r^C^r^-labeled NPSL2 RNAs. A portion of the 2D ^1^H-^1^H NOESY spectrum for A^2r^U^r^-labeled NPSL2 is shown in **Fig. 2B**. In this spectrum, inter-residue NOE cross peaks for all A-U base pairs were detected. For example, cross peaks were observed between A10.H2 and U34.H1’, and A16.H2 and U33.H1’, indicating that the A’s and U’s flanking the internal loop are base paired. Furthermore, the NOE cross peak observed between A11.H2 and U34.H1’ is consistent with A-form helical stacking of A11 on the preceding A10. Similarly, NOE cross peaks observed between A20.H2 and C30.H1’ in the 2D ^1^H-^1^H NOESY spectrum of A^2r^G^r^C^r^-labeled NPSL2 are also consistent with A-form helical stacking of C30 on C31 (**Fig. 2C**). The stretches of pyrimidine residues in the apical loop and upper helical region (residues 23-25 and 29-35) were significantly broader than those pyrimidines in the lower portion of the bottom stem. For these residues, sequential NOEs were obtained from analysis of ^1^H-^1^H NOESY spectra collected on A^8^C^H^- and A^8^U^6r^-labeled NPSL2 RNAs.

**Fig. 2.**
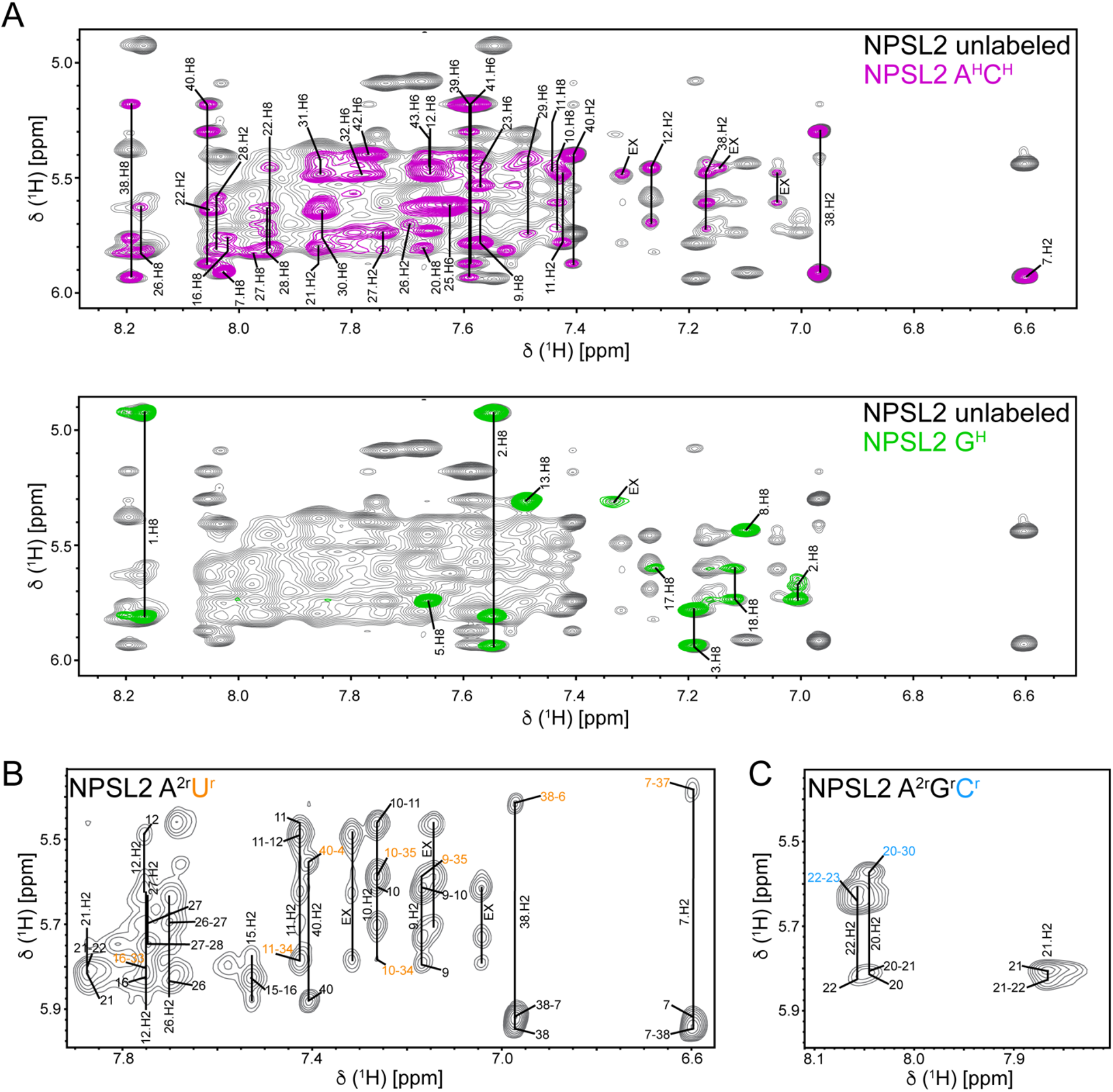
Deuterium-editing approach for NPSL2 chemical shift assignment. (A) Overlay of the ^1^H-^1^H NOESY spectra of unlabeled (black) and A^H^C^H^-labeled NPSL2 (magenta, top) and G^H^-labeled NPSL2 (green, bottom). (B) Region of the ^1^H-^1^H NOESY spectrum of A^2r^U^r^-labeled NPSL2, highlighting NOEs defining A.H2-U.H1’ NOEs (orange labels). (C) Region of the ^1^H-^1^H NOESY spectrum of A^2r^G^r^C^r^-labeled NPSL2 defining A.H2-C.H1’ NOEs in the apical loop (blue labels). EX denotes resonances in chemical exchange.

The 2D ^1^H-^1^H NOESY collected in 10% D_2_O was characterized by sharp and well-resolved peaks within the imino region (**Fig. S7**). We were therefore able to assign the imino proton signals for base pairs within A-helical regions of RNA (**Fig. S7**). Overall, 10 of the expected 14 base pairs of NPSL2 were unambiguously confirmed. However, the imino NOEs for G1 (from the terminal base pair) and the uracil residues flanking the internal loop (U33, U34, and U35) were not observed due to either overlap within the H1 and H3 signals or due to line broadening effects related to internal dynamics. While the imino protons for U33, U34, and U35 were not detectable from the imino walk, their base pairing was confirmed through the NOE analysis of the A^2r^U^r^-labeled NPSL2 sample (**Fig. 2B**).

Using the approaches outlined above, 100% of the non-exchangeable aromatic protons, 100% H1’, 100% H2’ and 100% H3’ protons were assigned (**Fig. S8A**), 71.4% imino protons were assigned (6 out of 7 base paired guanosine H1 and 3 out of 6 base paired uracil U-H3). Resonance assignments for NPSL2 have been deposited in the BMRB (NPSL2: 51350).

### Solution structure of the oncomiR-1 NPSL2 domain

The 3D structure of NPSL2 (**Fig. 3A**) was determined using a total of 413 NOE-derived ^1^H-^1^H distance restraints obtained from a combination of unlabeled and ^2^H-edited NOESY spectra (**Fig. S8B**). The base pairing was determined based on analysis of the 2D ^1^H-^1^H imino NOESY spectrum in 10% D2O and the 2D ^1^H-^1^H NOESY spectrum acquired on an A^2r^U^r^-labeled NPSL2 sample. The structure was gently refined with 27 ^1^H-^13^C residual dipolar couplings (RDCs) (**Fig. 3B**). The refined structure was in good agreement with the *ab initio* molecular reconstruction derived from small angle X-ray scattering (**Fig. 3A**). Guinier analysis was used to check for aggregation or repulsion and measure the radius of gyration (*R*_g_) from the Guinier fit (**Fig. S9**). The pair-distance distribution function [P(r)] curve (**Fig. 3C**) revealed the maximum distance (D_max_) to be 77 Å, consistent with the overall length of NPSL2. The most probable pair distance was ~20 Å, congruent with the averaged diameters of the A-helical stems, suggesting that NPSL2 adopts a largely A-helical geometry. Additionally, Kratky analysis suggests that NPSL2 is well folded, consistent with the derived hairpin structure (**Fig. S9, Table S1**). Furthermore, the back-calculated scattering data matched the experimental scattering data nicely, with a χ2 = 1.11 (**Fig. 3D**). A summary of NMR restraints and structural statistics are presented in **Table S2**.

**Fig. 3.**
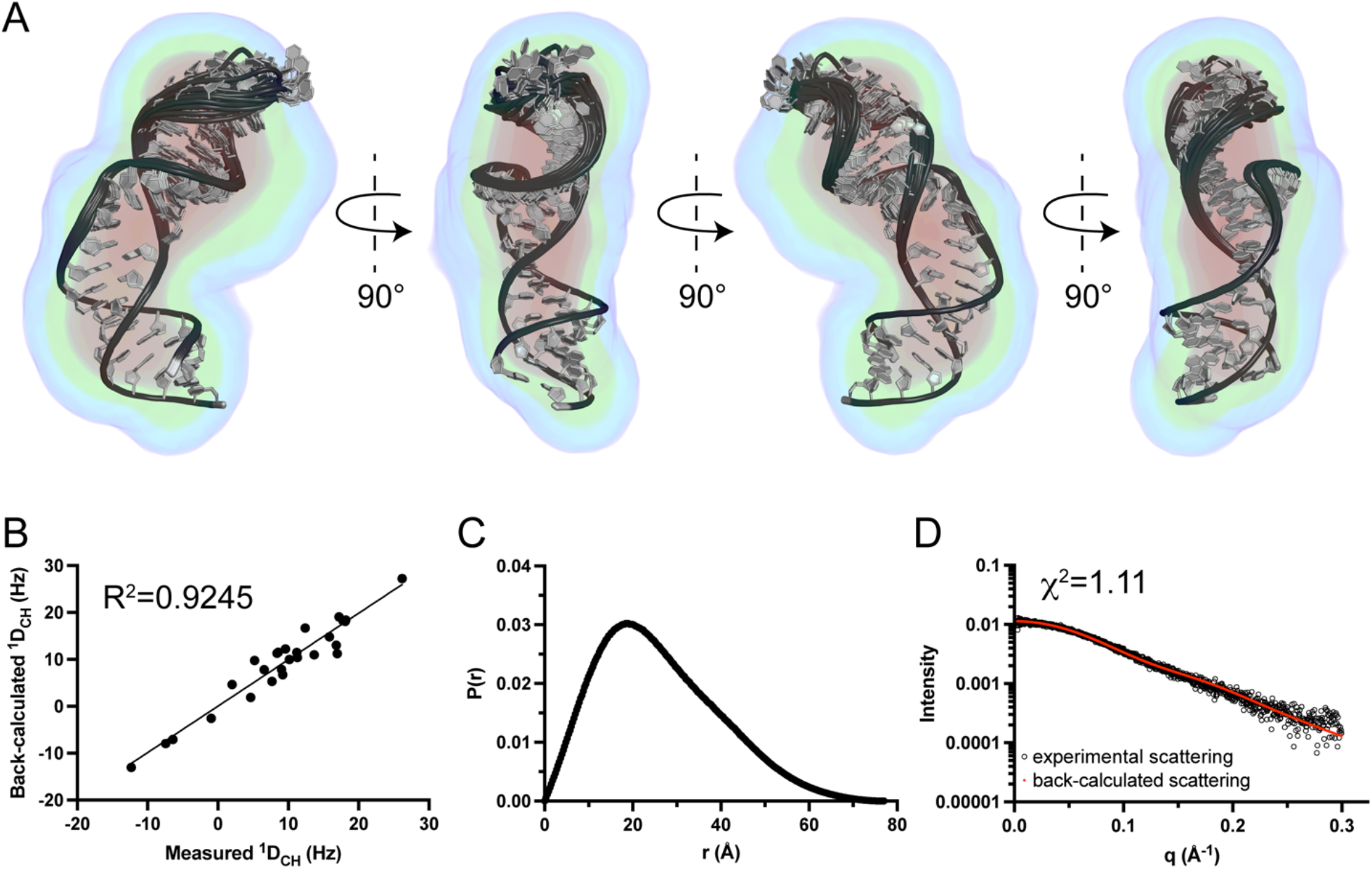
Solution structure of NPSL2. (A) Ensemble of 20 lowest energy AMBER structures aligned to the SAXS electron density map reconstruction. Contours of the density map are as follows: 15σ (red), 10σ (yellow), 7.5σ (green), 5σ (cyan), and 2.5σ (blue). (B) Correlation plot between measured and back-calculated RDCs for the lowest energy NPSL2 structure. (C) Pair distance distribution [P(r)] plot of NPSL2. (D) FoXS back-calculated scattering curve (red) of lowest energy structure fit to experimental SAXS data (black circles).

NPSL2 has large, but well-organized internal and apical loops. In the internal loop, A11 stacks on A10, extending the A-helical structure of the lower stem. We observed sequential aromatic-aromatic (i to i+1) and aromatic-anomeric (i to i-1) NOEs connecting A10, A11, A12, and G13, consistent with these bases being stacked. There is a break in the base stacking and significant kink in the phosphodiester backbone between G13 and U14, consistent with the lack of inter-residue NOEs between G13 and U14. A15 and U14 are stacked under A16, which is part of the upper helical segment (**Fig. 4A**).

**Fig. 4.**
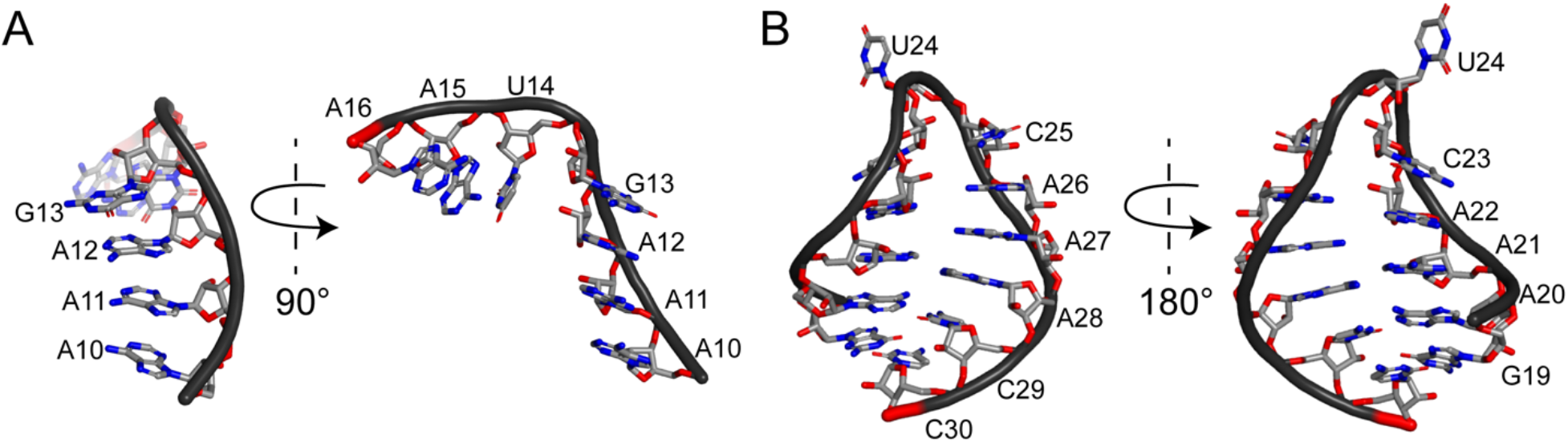
The internal and apical loops of NPSL2 are highly structured. (A) The NPSL2 internal loop. A11, A12, and G13 are stacked on A10 within the lower helical stem. There is a break in the helical stacking between G13 and U14 with a large kink in the phosphodiester backbone. U14 and A15 are staked beneath A16 within the upper helical stem. (B) The NPSL2 apical loop. A20-C23 are stacked directly on G19 and C25-C29 are stacked on C30, extending the A-helical structure of the upper helical stem. U24 is flipped out of the apical loop.

In the 10-nt apical loop, the first four nucleotides from the 5’-side of the loop (A20, A21, A22, and C23) extend the base stacking of the upper helical region, with A20 stacking directly on G19. The WC face of A20 is positioned toward the loop interior, with the possibility of forming a non-canonical base pair with C29. U24 is flipped out of the loop, consistent with break in inter-residue NOEs between U24 and the neighboring nucleotides. The stretch of nucleotides including C25, A26, A27, A28, and C29 form a continuous A-helical structure on the 3’-side of the loop with C29 directly stacking on C30 (**Fig. 4B**).

### A20 and C29 can form a stabilizing A^+^·C wobble pair at low pH

To probe the solution behavior of NPSL2, we performed CD thermal denaturation experiments under different pH conditions. The thermal stability of NPSL2 is enhanced upon lowering pH (**Fig. S10**). Protonation is a fundamental chemical property and the smallest modification on RNA and DNA molecules.^44,45^ The single stranded RNA and DNA nucleobases are typically uncharged, the intrinsic p*K*_a_ values for adenine and cytosine are far from neutrality.^46,47^ However, in some instances, the adenosine p*K*_a_ shifts towards neutrality, enabling their participation in non-canonical base pairings. The significance of pH-dependent base pairs has been demonstrated in a number of systems, including in miR processing,^44^ catalytic RNAs,^48–50^ viral RNAs.^51,52^

In the NPSL2 structure, A20 and C29 have the potential to form an A·C mismatch in the apical loop. A·C mismatches are stabilized at low pH, when the A.N1 is protonated, and can form an A^+^·C base pair.^46,49,53^ We performed a pH titration on ^13^C,^15^N-A-labeled NPSL2 and monitored chemical shift changes as a function of pH. Indeed, we observed chemical shift changes for A20.H8 upon reducing the pH (**Fig 5A**). A20 has an elevated p*K*_a_ value of 6.24 ± 0.24 (**Fig. 5A**), which is > 2.5 p*K*_a_ units greater than that of free AMP (p*K*_a_ = 3.5). As a control, we also followed the chemical shift changes of A7.H2 and A40.H2, which are present in the bottom stem. Consistent with their canonical base pairing, the chemical shifts of these bases are independent of pH (**Fig. S11**).

**Fig. 5.**
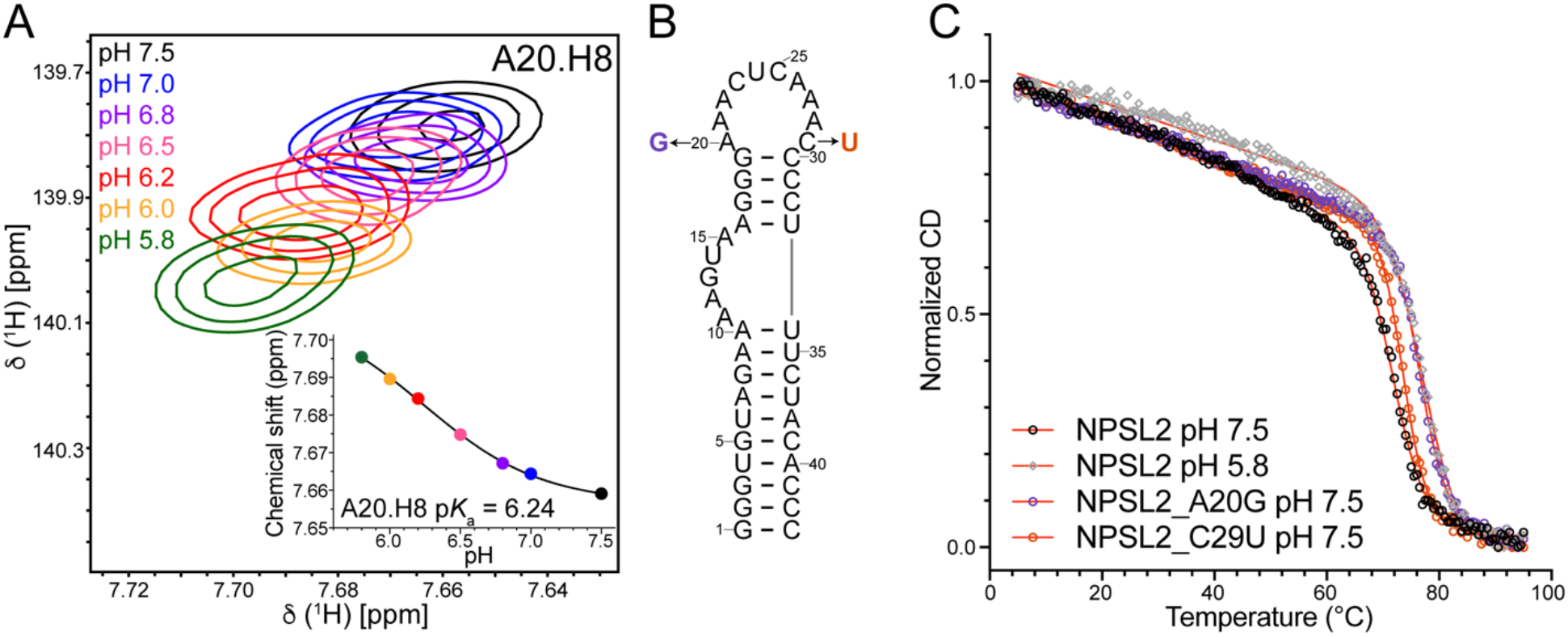
Protonation of A20 contribute stabilization of NPSL2. (A) pH dependence of the A20.H8 chemical shift. Plot of pH dependence of A20.H8 chemical shift is inset. (B) NPSL2 secondary structure highlighting single point mutations NPSL2_A20G and NPSL2_C29U. (C) Normalized CD thermal denaturation curves for NPSL2 at pH 7.5 (black), NPSL2 at pH 5.8 (grey), NPSL2_A20G at pH 7.5 (purple), and NPSL2_C29U at pH 7.5 (orange). Secondary structures were rendered using RNA2Drawer.^88^

To further probe the formation of an A20^+^·C29 base pair, we designed a single point substitution, NPSL2_C29U (**Fig. 5B**), to examine the stabilizing effect of canonical base pairing at this position. We expected NPSL2_C29U (pH 7.5) to be stabilized to a similar extent as NPSL2 (pH 5.8, where formation of the A20^+^·C29 base pair is promoted) due to the number of hydrogen bonds present in A-U and A^+^·C base pairs. However, we observed that NPSL2 (pH 5.8) was stabilized by 4.5 °C relative to NPSL2_C29U (pH 7.5) (**Fig. 5C**). We then made a second mutation, NPSL2_A20G (**Fig. 5B**), to compare the thermal stability. We found that NPSL2_A20G (pH 7.5) was slightly less stable (ΔT_m_=1.7 °C) than NPSL2 (pH 5.8) (**Fig. 5C** and **Table 1**). The additional stabilization of NPSL2 at low pH suggests that base pairing alone is not the sole source of the stabilizations, but rather there are regions outside of the A^+^·C mismatch that contribute to the pH-dependent stabilization of NPSL2.

**Table 1.** Thermodynamic properties for NPSL2 RNAs.

| RNA | $T_m^a$<br>(°C) | | $\Delta H^a$<br>(kcal/mol) | | $-\Delta S^b$<br>[cal/(mol*K)] | | $\Delta G^\circ$ (25°C)<br>(kcal/mol) | |
| --- | --- | --- | --- | --- | --- | --- | --- | --- |
|  | pH 7.5<sup>b</sup> | pH 5.8<sup>b</sup> | pH 7.5<sup>b</sup> | pH 5.8<sup>b</sup> | pH 7.5<sup>b</sup> | pH 5.8<sup>b</sup> | pH 7.5<sup>b</sup> | pH 5.8<sup>b</sup> |
| WT NPSL2 | 59.0 ± 0.3 | 63.9 ± 0.2 | -53 ± 1 | -43 ± 2 | 160 ± 4 | 130 ± 10 | -5.5 ± 0.2 | -5.0 ± 0.2 |
| NPSL2 | 72.0 ± 0.7 | 78.2 ± 0.3 | -89 ± 3 | -75 ± 6 | 260 ± 10 | 210 ± 20 | -12.2 ± 0.3 | -11.4 ± 1.0 |
| NPSL2_A20G | 76.5 ± 0.2 | 82.5 ± 0.2 | -104 ± 1 | -91 ± 3 | 300 ± 3 | 260 ± 10 | -15.4 ± 0.2 | -14.7 ± 0.5 |
| NPSL2_C29U | 73.7 ± 0.1 | 80.3 ± 0.1<sup>c</sup> | -122 ± 4 | -108 ± 3<sup>c</sup> | 350 ± 10 | 306 ± 5<sup>c</sup> | -17.1 ± 0.5 | -16.9 ± 0.3<sup>c</sup> |
| NPSL2_ΔAL | 74.9 ± 0.2 | 80.7 ± 0.1 | -100 ± 3 | -90 ± 2 | 290 ± 10 | 260 ± 10 | -14.3 ± 0.4 | -14.2 ± 0.3 |

<sup>a</sup> $T_m$ and $\Delta H$ values were obtained by fitting CD thermal denaturation profiles to a two-state model using sloping baselines.
<sup>b</sup> $\Delta S$ values are determined from **Equation 3**.
<sup>c</sup> Values and errors obtained from duplicate experiments.

### Protonation outside of A^+^·C mismatch contribute to stabilization of NPSL2

There is accumulating evidence that protonated nucleotides can stabilize RNA structure through electrostatic interactions.^54^ Both NPSL2_A20G and NPSL2_C29U, which remove the ionizable A^+^·C base pair, are further stabilized when lowering pH (**Fig. S12**). This observation is consistent with the idea that regions outside of the A^+^·C base pair contribute additional stabilization of NPSL2 at low pH. We next measured p*K*_a_ values of other adenosines in the apical loop by tracking their chemical shift changes as a function of pH. Most of the adenosines in the apical loop have overlapping chemical shifts and therefore could not be followed in a pH titration. We were able to measure the p*K*_a_ of A21 (based on detection of the A21.H2), which is significantly shifted towards neutrality (p*K*_a_ = 6.69) (**Fig. S13A**). Protonation of A21.N1 may stabilize the NPSL2 structure through formation of a cation-π interaction with the base of A22 (**Fig. S13B**). We were unable to measure the p*K*_a_ values of the remaining adenosines in the apical loop due to spectral overlap, however, protonation of A26, A27, and/or A28 might stabilize the NPSL2 apical loop through phosphate-base interactions (**Fig. S13C**).

To further isolate the effects of pH on NPSL2 stability, we generated an NPSL2 variant lacking the apical loop (NPSL2_ΔAL). The NPSL2_ΔAL RNA displayed increased thermal stability at low pH condition (**Fig. S14**). Therefore, the protonation of adenosines in the asymmetric internal loop may also contribute to the stabilization of NPSL2 at low pH. Here, we seek to understand this question via measuring the p*K*_a_ of other adenosines through pH titration. As can be seen in **Fig. S15**, all the adenosines in the internal loop showed elevated p*K*_a_ values. Based on the structure of the NPSL2 internal loop, protonation of A11, A12, and/or A15 can stabilize NPSL2 through forming cation-π interaction with A12, G13, and A16, respectively (**Fig. S15**).

RNA folding involves navigating the large conformational space of the polynucleotide chain. The favorable stacking exothermicities (enthalpy, ΔH) and the adverse conformational entropy (ΔS) are the predominate factors that determine the RNA folding stability.^55,56^ Based on our CD thermal denaturation experiments, NPSL2 displayed a higher T_m_ value at pH 5.8 relative to that at pH 7.5, indicating increased thermal stability (**Table 1**). It’s plausible that formation of an A20^+^·C29 wobble pair and favorable electrostatic interactions (cation-π interactions) at pH 5.8 make NPSL2 more stable. However, we observed the enthalpy at pH 5.8 is smaller than that at pH 7.5. Additionally, there is a smaller negative entropy change at pH 5.8 relative to pH 7.5. These enthalpic and entropic contributions result in similar, or slightly higher, free energy values for NPSL2 at pH 5.8 relative to pH 7.5(**Table 1**). Collectively, these findings indicate that NPSL2 upon thermal denaturation at pH 5.8, maintains partial base stacking. This is consistent with smaller negative entropy changes from the unfolded to folded state.

## Discussion

The precise control of oncomiR-1 processing is essential for normal development, and overexpression of certain miRs from this cluster is related to oncogenesis.^27^ Previous studies indicated that oncomiR-1 folded into a globular tertiary structure.^28,39–41^ However, there are no high-resolution structures of oncomiR-1 or its constituent pre-miRs, which greatly impacts our understanding of how RNA tertiary structure contributes to the regulation of oncomiR-1 processing.

Previous studies have shown that NPSL2 serves as an important element for regulating pri-miR-92a processing.^28,40,41^ However, how NPSL2 regulates pri-miR-92a processing remains unclear. In this study, we determined the three-dimensional structure of NPSL2 using a combination of solution NMR spectroscopy and SAXS (**Fig. 3**). This represents the first high-resolution structure determined of an oncomiR-1 RNA element. The internal and apical loops of NPSL2 are purine-rich and exhibit extensive base-stacking, giving rise to highly-ordered loop structures (**Fig. 4**). The 10-bp apical loop contains a pH-sensitive A^+^·C mismatch which is positioned at the top of the upper helical stem. Several adenosines within apical and internal loops have elevated p*K*_a_ values, which can further stabilize NPSL2 RNA structure (**Fig. 5** and **Fig. S15**).

Chaulk *et* al have shown that internal loop residues A11 and A12 are base stacked based on their cleavage by both ribonuclease (RNAse) 1 and RNAse V1.^41^ Similarly, they identified A20 and A21 in the apical loop as being stacked due to their RNAse 1 and V1 cleavage patterns. Both sets of findings are supported by our NMR data (**Fig.4 and Fig. S15**). Based on their RNAse cleavage assays, the base-pairing state of residues flanking the internal loop of NPSL2 could not be determined.^41^ However, in the NOESY spectrum of A^2r^U^r^-labeled NPSL2, we observed inter-residue NOE cross peaks between A10.H2 and U34.H1’, and A16.H2 and U33.H1’, indicating that the A’s and U’s flanking the internal loop are in fact base paired (**Fig.2B**). The NMR-derived structure therefore enriches our understanding of NPSL2 base pairing and stacking within the internal and apical loops of NPSL2.

Stretches of consecutive adenosines in single stranded regions, as are present in the NPSL2 internal and apical loops, are frequently involved in tertiary contacts that define the higher order structure of an RNA.^41^ Within the full-length oncomiR-1, the consecutive adenosines within the NPSL2 internal loop are solvent inaccessible, suggesting they are engaged in long-range RNA-RNA interactions.^28^ Site-specific photo-cross-linking was used to identify a long-range interaction between the single-stranded base-stacked adenosines in the internal loop of NPSL2 (A11, A12) with the stem of pre-miR-19b.^41^ Furthermore, the interaction between NPSL2 and pre-miR-19b serve as an RNA structure-based mechanism to suppress miR-92a processing.^41^

RNA binding proteins (RBPs) function as important post-transcriptional regulators of miR processing.^57,58^ For example, as a specific and posttranscriptional inhibitor of *let-7* biogenesis, Lin28 protein interacts with *let-7* precursor terminal loop and can inhibit both pri-*let-7* processing by Drosha^59,60^ and pre-*let-7* processing by Dicer.^61–63^ Within oncomiR-1, hnRNP-A1 serves as an auxiliary factor that promotes miR-18a biogenesis.^64^ KH-type splicing regulatory protein (KSRP) is known to bind pri-miR-20a and promote its processing.^57^ miR-92a is the least expressed oncomiR-1 derived miR *in vitro*,^40,41^ however, miR-92a is overexpressed in several tissues and tumor types.^65^ This suggests that specific RBPs may contribute to the structural remodeling within the 3’-core of oncomiR-1, exposing the pri-miR-92a cleavage sites. Therefore our future studies to characterize the structure-based regulation of pre-miR-92a expression are aimed at identifying RNA binding protein cofactors that regulate miR-92a processing.

## Material and methods

### Construct design and template preparation

Sequences of the RNAs examined in this study are listed in **Table S3**. The WT NPSL2 DNA template was designed with an upstream T7 promoter sequence and 5’-hammerhead ribozyme sequence to generate an RNA with a native 5’-UG starting sequence. The WT NPSL2 DNA template was amplified via polymerase chain reaction (PCR) using a set of four primers (**Table S4)** with EconoTaq PLUS 2x Master Mix (Lucigen). All the primers were ordered from Integrated DNA Technologies. The WT NPSL2_primer 4 contained 2’-*O*-methoxy modifications at the two 5’-most positions to reduce non-templated transcription.^66^ DNA oligonucleotides, which serve as templates for all other RNAs, were purchased from Integrated DNA Technologies (**Table S4**). The DNA oligonucleotides similarly contained 2’-*O*-methoxy modifications at the two 5’-most positions to reduce non-templated transcription.^66^ The DNA templates for *in vitro* transcription were generated by annealing the desired DNA oligonucleotide (40 μL, 200 μM) with an oligonucleotide corresponding to the T7 promoter sequence (20 μL, 600 μM) boiling for 3 min, and then slowly cooling down to room temperature. Then the annealed template was diluted with 940 μL H_2_O to produce the partially double-stranded DNA templates for *in vitro* transcription.

### RNA transcription

RNA transcription conditions were optimized prior to preparative scale transcription. RNAs were prepared by *in vitro* transcription in 1X transcription buffer (40 mM Tris base, 5 mM DTT, 1 mM spermidine, and 0.01% Triton-X, pH 8.5) with addition of 2-6 mM rNTPs (optimized for nucleotide content), 10-20 mM MgCl_2_, 8-13 pmol annealed or 1:10 (vol/vol) PCR amplified DNA template, 0.2 U/mL yeast inorganic pyrophosphatase (New England Biolabs), ~15 μM T7 RNA polymerase, 10-20% (vol/vol) DMSO. Reactions were incubated at 37 °C for 3-4 h and then quenched using a solution of 7 M urea and 500 mM EDTA (pH 8.5) and boiled for 3 min and snap cooled in ice water for 3 min. The transcription mixture was loaded onto preparative-scale 14% denaturing polyacrylamide gels for purification. Gels were run at constant power (20 W) for ~ 24 h for optimal resolution. The RNA product was visualized by UV shadowing and gel slices containing target RNA were excised. Gel slices were placed into an elutrap electroelution device (The Gel Company) in 1X TBE buffer. The RNA was eluted from the gel at constant voltage (120 V) for ~24 h. The eluted RNA was spin concentrated, washed with 2 M high-purity sodium chloride, and exchanged into water using Amicon-15 Centrifugal Filter Units (Millipore, Sigma). RNA purity was checked by running RNA on a 14% analytical denaturing gel. RNA concentration was quantified via UV-Vis absorbance.

Isotopically-labeled RNAs were produced as described above by replacing the rNTP mixture with rNTPs of appropriate isotope labeling. The partially- and per-deuterated rNTPs used for *in vitro* transcription were obtained from Cambridge Isotope Laboratories (CIL, Andover, MA) or generated in house, as described below. ^15^N/^13^C rNTPs were obtained from Cambridge Isotope Laboratories (CIL, Andover, MA)

### Selective C8 protiation and deuteration of ATP and GTP

Protiation at the C8 position of perdeuterated rGTP and rATP was achieved by incubation with triethylamine (TEA, 5 equiv) in H_2_O (60 °C for 24 h and for 5 days, respectively). Deuteration of the C8 position of fully protiated GTP and ATP was achieved by analogous treatment with D_2_O (99.8% deuteration; CIL). TEA was subsequently removed by lyophilization.

### Native gel electrophoresis

RNA (20 μM) was prepared in 50 mM potassium phosphate, pH 7.5, 1mM MgCl_2_ and incubated at 4 °C for 30 min. Then 20% (vol/vol) 50% glycerol was added to the RNA sample. The sample was immediately mixed and loaded onto 16% native polyacrylamide gel. The gel was run at 100 V at room temperature. The native gel and running buffer were prepared to a final 1X buffer of 89 mM Tris base, 89 mM boric acid (pH ~8.3) to better preserve the RNA conformation. ^67^

### Size-exclusion chromatography

Chromatographic separation was performed with NGC™ Chromatography system (Bio-Rad) with a UV detection at 260 nm. The conformational homogeneity of RNA sample was checked with an ENrich™ SEC 650 10 x 300 Column (Bio-Rad; column volume: 24 mL). RNA (20 μM, 100 μL) in 50 mM potassium phosphate, pH 7.5 was boiled 3 min, snap cooled 3 min, then MgCl_2_ were added to a final concentration of 1 mM. The sample was incubated 30 min before loading onto the column. A flow rate of 1 mL/min was used in all the chromatographic steps.

### NMR data acquisition, processing, and analysis

NMR spectra were collected on 600 and 800 MHz Bruker AVANCE NEO spectrometers equipped with a 5 mm TCI cryogenic probe (University of Michigan BioNMR Core) (see **Table S5** for experimental parameters). NMR data were processed with NMRFx^68^ and NMRPipe^69^ and analyzed with NMRViewJ^70^ and MestReNova. ^1^H chemical shifts were referenced to water and ^13^C chemical shifts were indirectly referenced from the ^1^H chemical shift.^71^

^1^H-^13^C residual dipolar couplings (RDCs) were carried out by measurement of ^1^H-^13^C doublet splittings. Filtering of the downfield and upfield doublet components into separate spectra via the inphase-antiphase (IPAP) HSQC experiments^72–74^ under both isotropic and anisotropic (Pf1 bacteriophage at ~ 12.5 mg/mL, ASLA) conditions with a selective ^13^C/^15^N-A/G-labeled NPSL2 (fully protonated C and U) in 50 mM potassium phosphate buffer and 1 mM MgCl_2_, pH 7.5 at 15 °C. Phage concentration was verified via ^2^H splitting (~10.0 Hz) at 800 MHz. Residual dipolar coupling values were determined by taking difference in ^1^J_CH_ couplings under anisotropic and isotropic conditions.

### pH titration and p*K*_a_ measurements

Changes in chemical shift as a function of pH were monitored by SOFAST-HSQC experiments^75^ using selective ^13^C/^15^N-A-labelled (fully protonated C, G and U) NPSL2 samples. Here, the ^13^C/^15^N-A-NPSL2 was buffer exchanged into water, lyophilized, and then resuspended in buffer of the appropriate pH (50 mM potassium phosphate, 1 mM MgCl_2_, pH =5.8, 6.0, 6.2, 6.5, 6.8, 7.0, 7.3 or 7.5). SOFAST-HSQC spectra were acquired at 15 °C. The changes in ^1^H or ^13^C chemical shifts were followed throughout the pH titration. The curves of chemical shift changes versus pH values were used to determine p*K*_a_ values (**Equation 1**):^46,47^

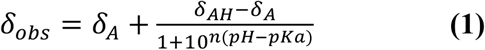

where δ_A_ is the chemical shift at high pH, δ_AH_ is the chemical shift at low pH, and δ_obs_ is the observed chemical shift at a given pH, and n is the Hill coefficient, which may be related to the number of ionization events.

### CD-thermal denaturation of RNA and data analysis

CD-thermal denaturation experiments were performed on JASCO J-1500 CD spectrometer with a heating rate of 1 °C per min between 5 °C to 95 °C. Data points were collected every 0.5 °C with absorbance detection at 260 nm. RNA samples (20 μM) were prepared in 50 mM potassium phosphate buffer (pH 7.5, 6.5, 5.8, or 5.45) with 1 mM MgCl_2_. The obtained reversible single-transition melting profiles were analyzed by a two-state model using sloping baselines (**Equation 2**): ^76^

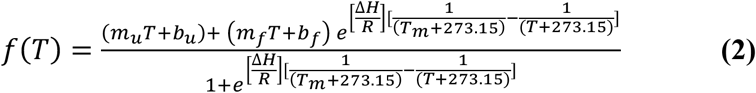

where m_u_ and mf are the slopes of the lower (unfolded) and upper (folded) baselines, b_u_ and b_f_ are the y-intercepts of the lower and upper baselines, respectively. ΔH (in kcal/mol) is the enthalpy of folding and T_m_ (in °C) is the melting temperature, R is the gas constant (0.001987 kcal/(Kmol)). Experiments were performed in triplicate or duplicates. The entropy change (ΔS) was determined by **Equation 3**:

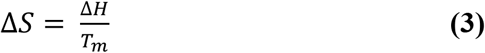

The standard free energy change (ΔG°) was calculated at 25 °C using ΔH and ΔS. Experiments were performed in triplicate and were in good agreement.

### SEC-SAXS data acquisition and analysis

SAXS was performed at BioCAT (beamline 18ID at the Advanced Photon Source, Chicago) in 50 mM potassium phosphate buffer, pH 7.5, 1 mM MgCl_2_, and 50 mM NaCl at 20 °C, with in-line size exclusion chromatography (SEC) to separate the homogeneous RNA sample from aggregates and other contaminants, ensuring optimal sample quality. NPSL2 (250 μL, 3.8 mg/mL) was loaded onto a Superdex 75 Increase 10/300 GL column (Cytiva), and run at 0.6 mL/min by an AKTA Pure FPLC (GE) with UV monitoring at 260 nm. The eluate was passed through the 1.0 mm ID quartz capillary SAXS flow cell with ~20 μm walls. A coflowing buffer sheath was used to separate the sample from the capillary walls, reducing radiation damage.^77^ Scattering intensity was recorded using an Eiger2 XE 9M (Dectris) detector which was placed 3.678 m from the sample giving us access to a q-range of 0.0028 Å^-1^ to 0.4157 Å^-1^. 0.5 s exposures were acquired every 1 s during elution and data was reduced using BioXTAS RAW 2.1.1.^78^ Buffer blanks were created by averaging regions preceding the elution peak and subtracted from exposures selected from the elution peak to create the *I*(*q*) vs *q* curves used for subsequent analyses. The GNOM package was used to determine the pair-distance distribution function [*P*(r)] required for molecular reconstruction.^79^ The maximum linear dimension of the molecule, *D*_max_, is calibrated for goodness-of-fit by enforcing a smooth zeroing of *P*(*D*_max_). DENSS was used to calculate the *ab initio* electron density map directly from the GNOM output.^80^ 20 reconstructions were performed in slow mode using default parameters and averaged. Alignment of the reconstruction to the structure was achieved using the DENSS alignment function in BioXTAS RAW. Reconstructions were visualized using PyMOL 2.5.2. FoXS was used to back-calculate the scattering profile of NPSL2.^81,82^

### Structure calculations

Initial structures were generated with CYANA simulated annealing molecular dynamics calculations over 128,000 steps. Upper limits for the NOE distance restraints were binned based on peak intensity and were generally set at 5.0 Å for weak, 3.3 Å for medium, and 2.7 Å for strong signals. Exceptions included intraresidue NOEs between H6/H8 and H2’ (4.0 Å) and H3’ (3.0 Å). 6.0 Å upper limit restraints were used for very weak signals, including sequential H1’-H1’ NOEs and intraresidue H5-H1’ NOEs. Standard torsion angle restraints were used for regions with A-helical geometry, allowing for ± 25° deviations from ideality (ζ=-73°, α=-62°, β=180°, γ=48°, δ=83°, ε=-152°). Standard hydrogen bonding and planarity restraints were used, and cross-strand P–P distance restraints were employed for A-form helical regions to prevent the generation of structures with collapsed major grooves.^83^ A grid search was performed over a broad range of tensor magnitude and rhombicity with weighting of the experimentally determined ^1^H-^13^C residual dipolar couplings (RDCs) constraints.

The 20 CYANA-derived structures were then subjected to molecular dynamics simulations and energy minimization with AMBER.^84^ The upper limit NOE, hydrogen bond, and dipolar coupling restraints were used, together with restraints to enforce planarity of aromatic residues and standard atomic covalent geometries and chiralities.^83,85^ Backbone torsion angle and phosphate-phosphate restraints were not used during AMBER refinement. Calculations were performed using the RNA.OL3^86^ and generalized Born^87^ force fields.

## Acknowledgements

We thank Dr. Jesse Hopkins for assistance with SAXS data collection and processing. This work was supported by National Institute of General Medical Sciences of the National Institutes of Health grant R35 GM138279 (S.C.K.), the Pew Charitable Trusts Scholars Program (S.C.K.), the National Institutes of General Medical Sciences training grant T32 GM132046 (A.M.), and a Rackham Merit Fellowship from the University of Michigan (A.M.). Research reported in this publication was supported by the University of Michigan BioNMR Core Facility (U-M BioNMR). U-M BioNMR Core is grateful for support from U-M including the College of Literature, Sciences and Arts, Life Sciences Institute, College of Pharmacy and the Medical School along with the U-M Biosciences Initiative. This research used resources of the Advanced Photon Source, a U.S. Department of Energy (DOE) Office of Science User Facility operated for the DOE Office of Science by Argonne National Laboratory under Contract No. DE-AC02-06CH11357. This project was supported by grant P30 GM138395 from the National Institute of General Medical Sciences of the National Institutes of Health. Use of the Pilatus 3 1M detector was provided by grant 1S10OD018090-01 from NIGMS. The content is solely the responsibility of the authors and does not necessarily reflect the official views of the National Institute of General Medical Sciences or the National Institutes of Health.

## Data availability

Resonance assignments have been deposited in the BMRB (NPSL2_Frag1: **<u>51348</u>**, NPSL2_Frag2: **<u>51349</u>**, NPSL2: **<u>51350</u>**). NMR-derived structures have been deposited in the PDB (NPSL2: **<u>7UGA</u>**). Experimental SAXS data of NPSL2 has been deposited in the Small Angle Scattering Biological Data Bank under accession code **<u>SASDNJ7</u>**.

