## Supplemental Information for "Solution structure of NPSL2, a regulatory element in the oncomiR-1 RNA"

**Table S1.** SEC-SAXS data acquisition, sample details, data analysis, model fitting, and software used.

| | |
|---|---|
| **(*a*) Sample details** | |
| Sample | NPSL2 |
| Organism | *Homo sapiens* |
| Sequence (5′→3′) | GGGUGUAGAAAAGUAAGGGAAACUCAAACCC CUUUCUACACCC |
| Extinction coefficient $\varepsilon_{280}$ ($M^{-1}cm^{-1}$) | 415000 |
| Molecular mass *M* from chemical composition (Da) | 13850 |
| Loading volume (μl) | 250 |
| Concentration (mg/ml) | 3.8 |
| Flow rate (ml/min) | 0.6 |
| Solvent composition | 50 mM potassium phosphate buffer (pH = 7.5), 1 mM $MgCl_2$, 50 mM NaCl |
| **(*b*) SAS data collection parameters** | |
| Instrument | BioCAT facility at the Advanced Photon Source beamline 18ID with Pilatus3 X 1M (Dectris) detector |
| Wavelength (Å) | 1.033 |
| Beam size ($\mu m^2$) | 150 (h) x 25 (v) focused at the detector |
| Camera length (m) | 3.678 |
| *q*-measurement ($Å^{-1}$) | 0.0028 to 0.42 |
| Absolute scaling method | Glassy Carbon, NIST SRM 3600 |
| Basis for normalization to constant counts | To transmitted intensity by beam-stop counter |
| Method for monitoring radiation damage | Automated frame-by-frame comparison of relevant regions using CORMAP[1] implemented in BioXTAS RAW |
| Exposure time, number of exposures | 0.5 s exposure time with a 1 s total exposure period (0.5 s on, 0.5 s off) of entire SEC elution |
| Sample configuration | SEC-SAXS with sheath-flow cell,[2] effective path length 0.542 mm. Size separation by an AKTA Pure with a Superdex 75 Increase 10/300 GL column. |
| Sample temperature (°C) | 20 |
| **(*c*) Software employed for SAS data reduction, analysis, and interpretation** | |
| SAXS data reduction | Radial averaging, frame comparison, averaging, and subtraction done using BioXTAS RAW 2.1.1.[3] |
| Basic analysis: Guinier, M.W., P(r), Shape modelling | Guinier fit and M.W. using BioXTAS RAW, P(r) using GNOM.[4] RAW uses MoW and Vc M.W. methods.[5,6] Density map reconstructions using DENSS.[7] |
| Molecular graphics | PyMOL 2.5.2 |

| | |
|---|---|
| **(*d*) Structural parameters** | |
| Guinier Analysis | |
| $I(0)$ ($cm^{-1}$) | 0.0111 (Å) 3.79e-5 |
| $R_g$ (Å) | 19.7018 ± 0.19 |
| $q$-range ($Å^{-1}$) | 0.0034 to 0.05094 |
| Quality-of-fit ($r^2$) | 0.856 |
| *P*(*r*) analysis | |
| $I(0)$ ($cm^{-1}$) | 0.01 ± 2.79e-5 |
| $R_g$ (Å) | 20.78 ± 0.09 |
| $d_{max}$ (Å) | 77.0 |
| $q$-range ($Å^{-1}$) | 0.0034 to 0.3002 |
| Quality-of-fit ($\chi^2$) | 1.084 |
| Porod Volume ($Å^3$) | 14100 |
| **(*e*) Shape modelling results** | |
| *DENSS* (default parameters, 20 calculations) | |
| $q$-range for fitting | 0.0034 to 0.3002 |
| Symmetry, anisotropy assumptions | P1, none |
| Ambiguity score (AMBIMETER) | |
| $\chi^2$ range | 2.644 |
| Model resolution (Å) | 0.0014 to 0.01252 |
| | 21.8 |
| **(*f*) Atomistic modelling** | |
| NMR structures | PDB entry 7UGA |
| *FoXS* | |
| $\chi^2$ | 1.11 |
| Predicted $R_g$ (Å) | 20.41 |
| $c_1$, $c_2$ | 1.02, 3.64 |
| **(*g*) SASBDB IDs for data and models** | |
| NPSL2 | SASDNJ7 |

**Table S2.** NMR Restraints and Structure Statistics[1]

| Protein Data Bank accession code | 7UGA |
|---|---|
| Cyana[2] | |
| NOE-derived restraints | 413 |
| Intraresidue | 86 |
| Sequential | 242 |
| Long range (\|i – j\| > 1) | 14 |
| H-bond restraints | 72 |
| NOE restraints/residue | 9.6 |
| Total dihedral angle restraints | 226 |
| RDC restraints | 27 |
| Q-factor (%) | 17.29 ± 0.43 |
| Target function (Å$^2$) | 0.30 ± 0.0054 |
| Amber[3] | |
| Amber energy | -9874.48 |
| Distance | 95.58 |
| Torsion | 3.47 |
| RMSD (Å)[4] | 1.24 ± 0.35 |
| RMSD (1-10, 34-43) (Å)[4] | 0.228 ± 0.092 |
| MolProbity analysis[5] | |
| Clashscore | 0 ± 0 |
| Probably wrong sugar pucker (%) | 0 ± 0 |
| Bad backbone conformation (%) | 4.19 ± 1.21 |
| Bad bonds (%) | 0 ± 0 |
| Bad angles (%) | 0 ± 0 |

[1] Statistics were calculated over the entire structure (residues G1-C43) unless otherwise specified.
[2] Statistics for the 20 structures with lowest target function.
[3] Statistics for the 20 lowest energy structures.
[4] Mean ± standard deviation for all heavy atoms, over the residues listed in parentheses.
[5] The 20 amber-refined structures were evaluated using the MolProbity webserver.[8]

**Table S3.** RNA constructs.

| Construct | RNA sequence 5'→3' [a] |
|---|---|
| WT NPSl2 | UGUAGAAAAGUAAGGGAAACUCAAACCCCUUUCUACA |
| NPSL2 | GGGUGUAGAAAAGUAAGGGAAACUCAAACCCCUUUCUACACCC |
| NPSL2_frag1 | GGGUGUAGAAGAGAUUCUACACCC |
| NPSL2_frag2 | GGAGGGAAACUCAAACCCCUCC |
| NPSL2_ΔAL | GGGUGUAGAAAAGUAAGGGGAGACCCUUUCUACACCC |
| NPSL2_A20G | GGGUGUAGAAAAGUAAGGGGAACUCAAACCCCUUUCUACACCC |
| NPSL2_C29U | GGGUGUAGAAAAGUAAGGGAAACUCAAAUCCCUUUCUACACCC |

[a] Red nucleotides indicate non-native nucleotides corresponding to the GAGA tetraloop sequence.

**Table S4.** RNA primers.

| Primer | Primer sequence 5'→3' [a] |
|---|---|
| WT_NPSL2_primer 1 | TTCTAATACGACTCACTATATAATACGACTCACTATAGGGATCACTTTTCTACACTGATG |
| WT_NPSL2_primer 2 | CCGGGTACCGTTTCGTCCTCACGGACTCATCAGTGTAGAAAAGTGATCCCTA |
| WT_NPSL2_primer 3 | CGAAACGGTACCCGGTACCGTCTGTAGAAAAGTAAGGGAAACTCAA |
| WT_NPSL2_primer 4 | mUmGTAGAAAGGGGTTTGAGTTTCCCTTACTTTTCTACAGA |
| NPSL2 | mGmGGTGTAGAAAGGGGTTTGAGTTTCCCTTACTTTTCTACACCCTATAGTGAGTCGTATTA |
| NPSL2_Frag1 | mGmGGTGTAGAATCTCTTCTACACCCTATAGTGAGTCGTATTA |
| NPSL2_Frag2 | mGmGAGGGGTTTGAGTTTCCCTCCTATAGTGAGTCGTATTA |
| NPSL2_ΔAL | mGmGGTGTAGAAAGGGTCTCCCCTTACTTTTCTACACCCTATAGTGAGTCGTATTA |
| NPSL2_A20G | mGmGGTGTAGAAAGGGGTTTGAGTTCCCCTTACTTTTCTACACCCTATAGTGAGTCGTATTA |
| NPSL2_C29U | mGmGGTGTAGAAAGGGATTTGAGTTTCCCTTACTTTTCTACACCCTATAGTGAGTCGTATTA |
| T7 promoter | TAATACGACTCACTATA |

[a] m denotes 2'-O-Me modification of the primer

**Table S5.** NMR experimental parameters.

| **Experimental parameter** | **$^1$H-$^1$H NOESY (100% $D_2O$ or 90% $H_2O$/10% $D_2O$)** | **$^1$H-$^1$H TOCSY** | **$^1$H-$^{13}$C HMQC** | **SOFAST-HMQC** | **$^1$H-$^{13}$C IPAP-HSQC (isotropic)** | **$^1$H-$^{13}$C IPAP-HSQC (Pf1 aligned; anisotropic)** |
|---|---|---|---|---|---|---|
| **pulse sequence** | noesyesgpph[9] | mlevphpr.2[10] | hmqcphpr[11] | sfhmqc.aro[12] | hsqcetgpiajcsp.2[13] | hsqcetgpiajcsp.2[13] |
| **ds** | 16 | 16 | 16 | 16 | 16 | 16 |
| **ns** | 24 | 24 | 336 | 24 | 96 | 152 |
| **sw(F2)** | 9.8104 | 9.8104 | 9.8104 | 8.7777 | 20.1549 | 20.1549 |
| **sw(F1)** | 9.8104 | 9.8104 | 50 | 10 | 40 | 40 |
| **TD(F2)** | 8192 | 2048 | 1058 | 8192 | 16384 | 16384 |
| **TD(F1)** | 800 | 600 | 88 | 512 | 256 | 256 |
| **O1 (ppm)** | 4.703 | 4.703 | 4.703 | 4.700 | 4.703 | 4.703 |
| **O2 (ppm)** | - | - | 148.00 | 139 | 142 | 142 |
| **D1 (s)** | 3 | 2 | 1.5 | 0.2 | 1 | 1 |
| **mixing time (ms)** | 300 | 80 | - | - | - | - |
| **experiment time (h)** | ~27 | ~8 | ~12 | ~4 | ~21 | ~33 |

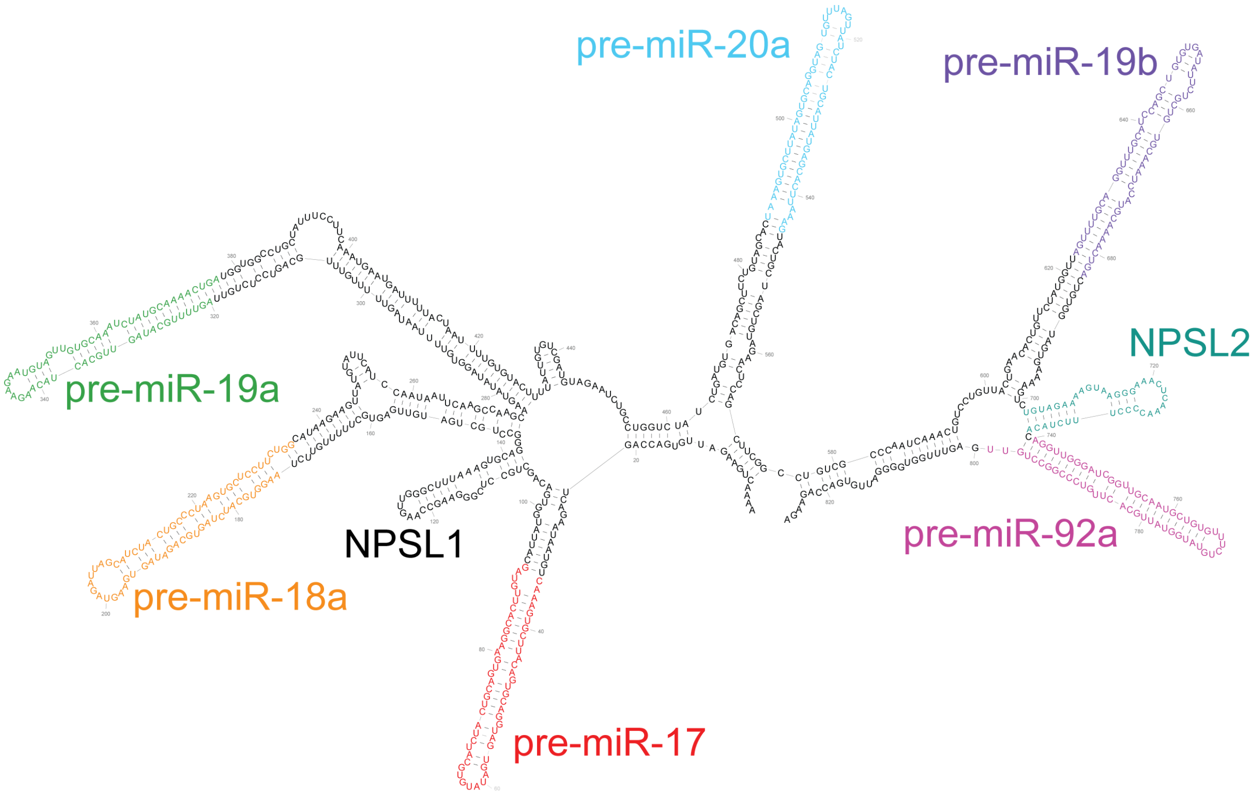


**Figure S1**

**Figure S1. Secondary structure of oncomiR-1.** Secondary structure derived from chemical probing.[14] Secondary structures were rendered using RNA2Drawer.[15]

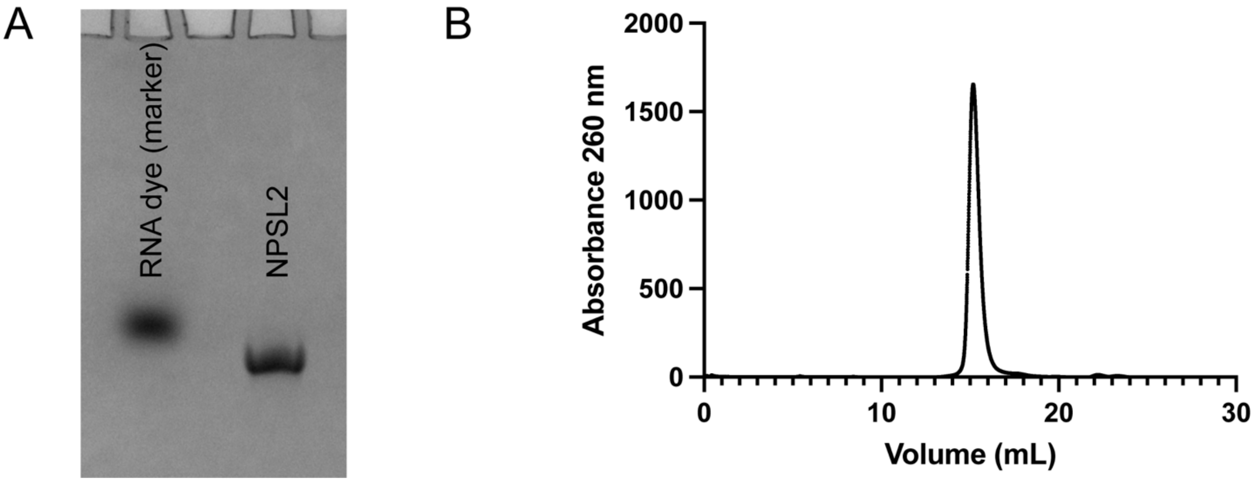


**Figure S2**

**Figure S2. NPSL2 is homogeneous in solution.** (A) Native gel electrophoresis of NPSL2 RNA. (B) Size exclusion chromatography of NPSL2 RNA.

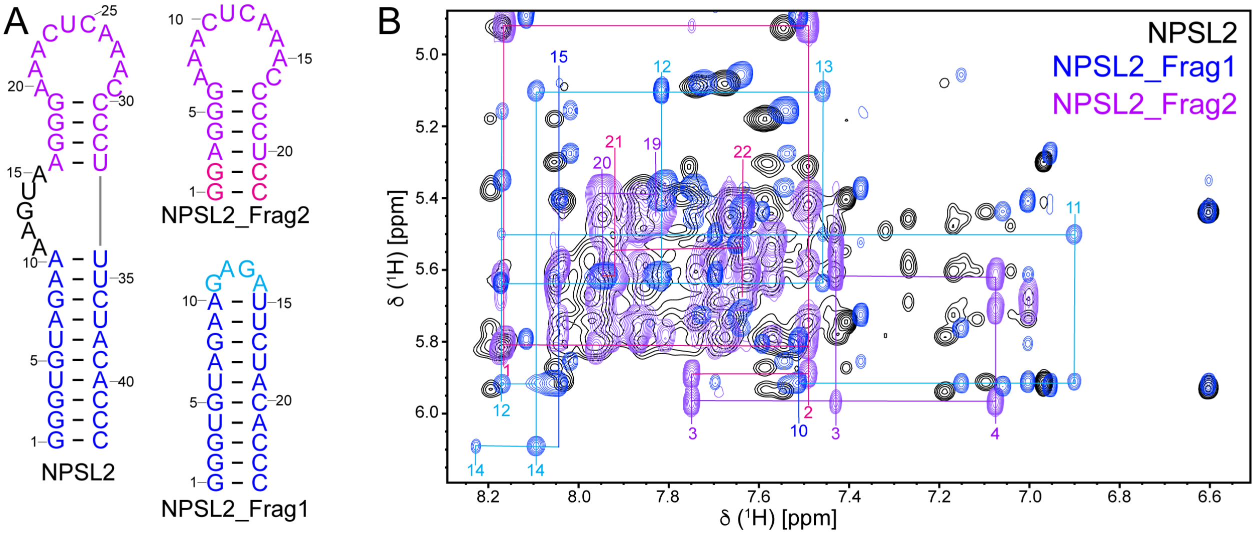


**Figure S3**

**Figure S3. Oligonucleotide-based approach for NPSL2.** (A) Secondary structure of NPSL2, NPSL2_Frag1, and NPSL2_Frag2. NPSL2_Frag1corresponds to the bottom stem of NPSL2 and capped with a GAGA tetraloop (light blue). NPSL2_Frag2 contains the upper stem of NPSL2 and has two non-native G-C base pairs to stabilize the hairpin (pink). (B) Overlay of the $^1$H-$^1$H NOESY spectra of NPSL2 (black), NPSL2_Frag1 (blue), and NPSL2_Frag2 (purple). Select resonance assignments are shown for NPSL2_Frag1, and NPSL2_Frag2. Colors and numbering are as indicated in (A). Secondary structures were rendered using RNA2Drawer.[15]

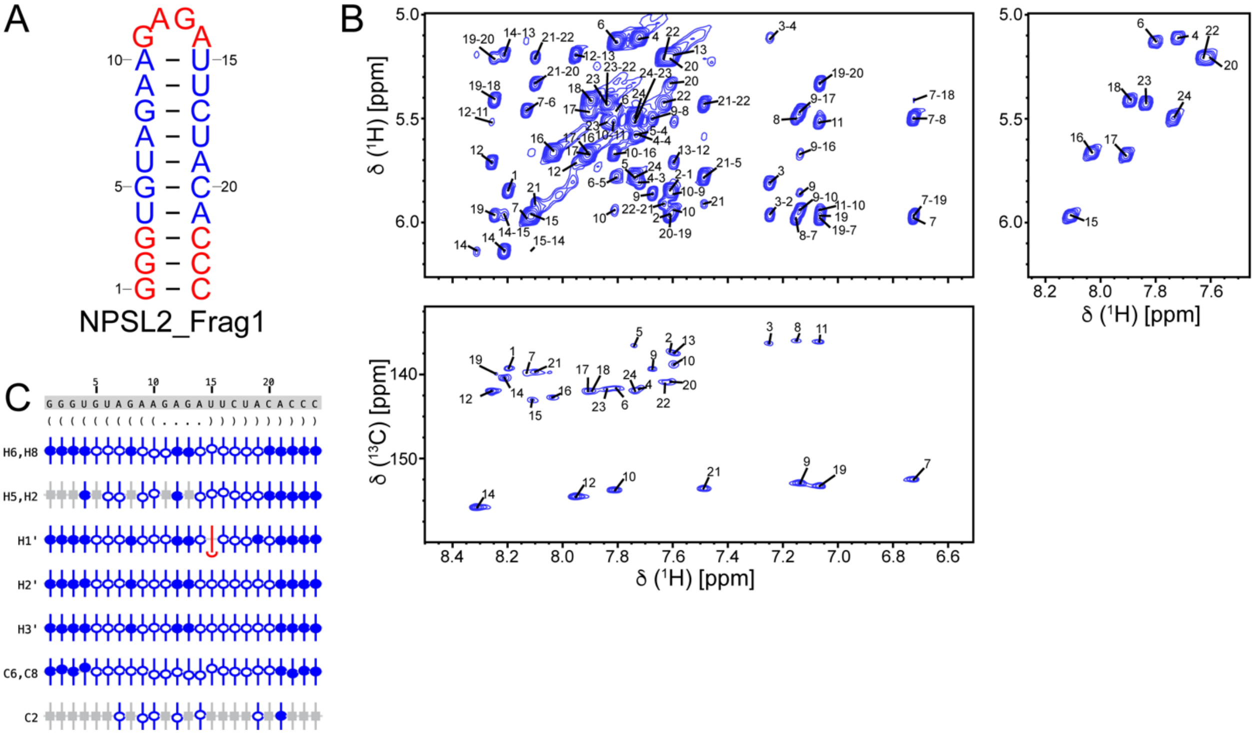


**Figure S4**

**Figure S4. Assigned chemical shifts of NPSL2_Frag1 RNA.** (A) The secondary structure of NPSL2_Frag1. (B) $^1$H-$^1$H NOESY (top, left), $^1$H-$^1$H TOCSY (top, right) and $^1$H-$^{13}$C HMQC (bottom). (C) Summary of the sequence, secondary structure, and assignment validation for NPSL2_Frag1. The secondary structure is shown in dot bracket notation. Assigned chemical shifts (H6/H8, H5/H2, H1′, H2′, H3′, C6/C8, C2) are indicated with open and filled circles. The vertical offset of the circles indicates the deviation from the predicted values for that atom. Filled circles indicate that there are chemical shifts for atoms with the same set of attributes in the BMRB. Open circles indicate atoms that have a prediction, but for which no exact matches of the attributes are available in the BMRB. Grey boxes represent atoms that are not present in a given base. Secondary structures were rendered using RNA2Drawer.[15]

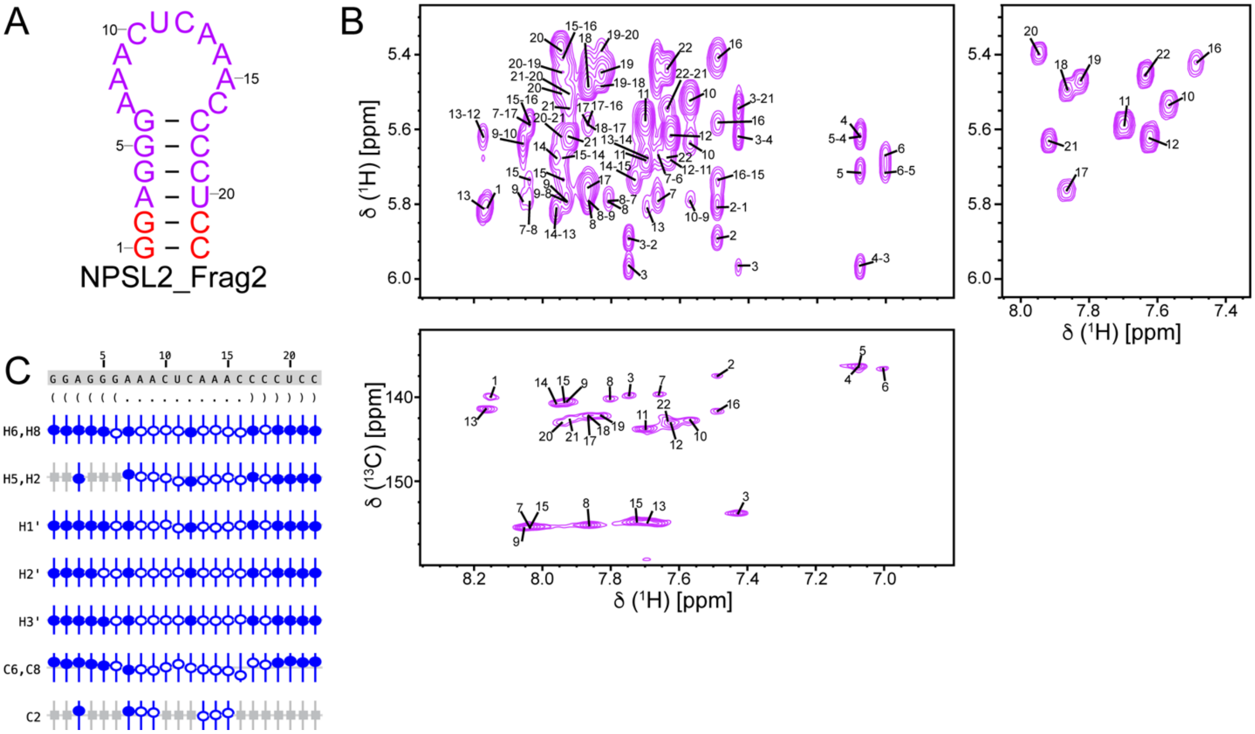


**Figure S5**

**Figure S5. Assigned chemical shifts of NPSL2_Frag2 RNA.** (A) The secondary structure of NPSL2_Frag2. (B) $^{1}H$-$^{1}H$ NOESY (top, left), $^{1}H$-$^{1}H$ TOCSY (top, right) and $^{1}H$-$^{13}C$ HMQC (bottom). (C) Summary of the sequence, secondary structure, and assignment validation for NPSL2_Frag2. The secondary structure is shown in dot bracket notation. Assigned chemical shifts (H6/H8, H5/H2, H1′, H2′, H3′, C6/C8, C2) are indicated with open and filled circles. The vertical offset of the circles indicates the deviation from the predicted values for that atom. Filled circles indicate that there are chemical shifts for atoms with the same set of attributes in the BMRB. Open circles indicate atoms that have a prediction, but for which no exact matches of the attributes are available in the BMRB. Grey boxes represent atoms that are not present in a given base. Secondary structures were rendered using RNA2Drawer.[15]

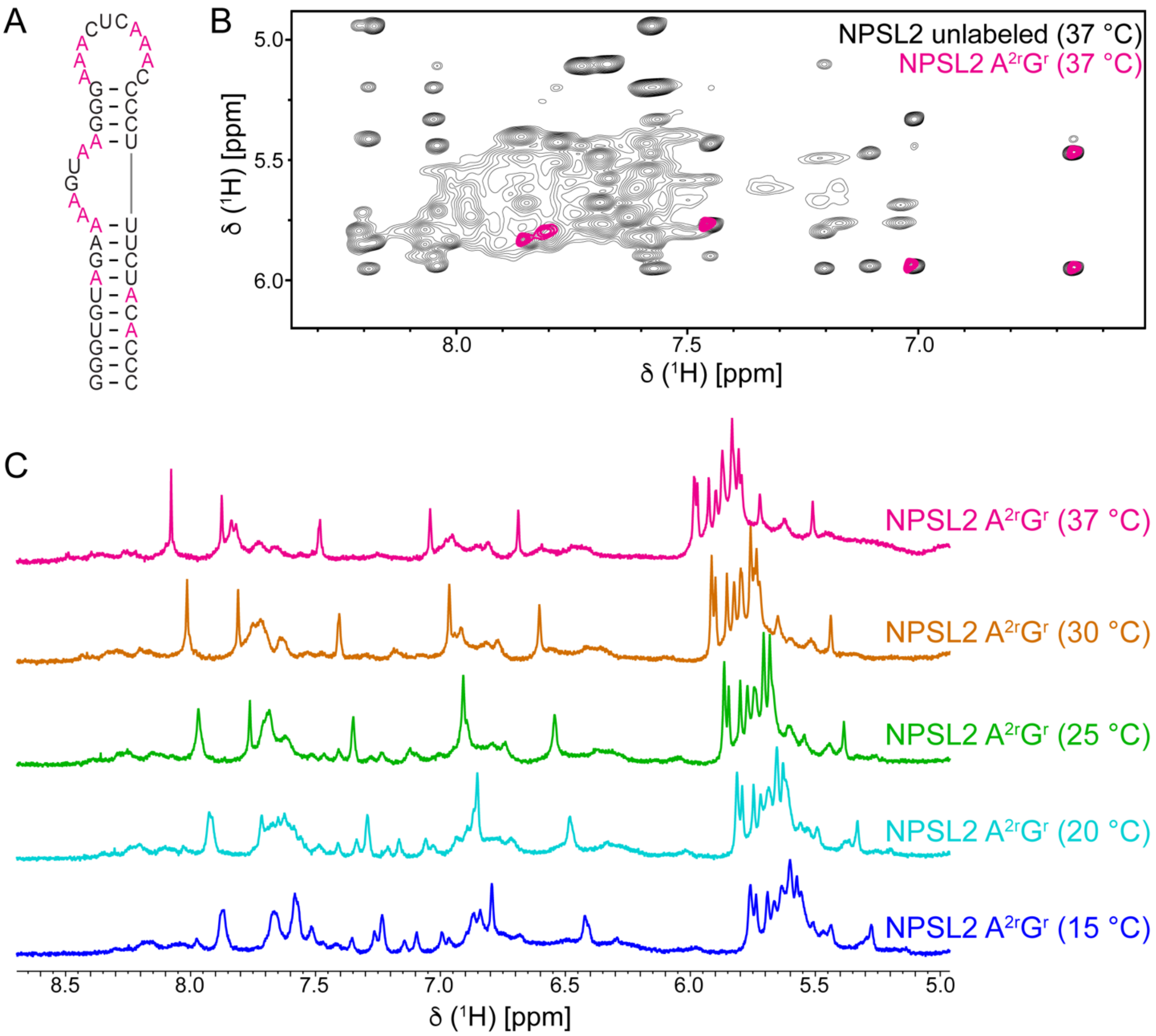


**Figure S6**

**Figure S6. Temperature-dependence of NPSL2 NMR spectrum.** (A) Secondary structure of NPSL2. All adenosines in the sequence are colored pink. (B) $^{1}H$-$^{1}H$ NOESY spectrum of unlabeled NPSL2 (black) overlaid with that of $A^{2r}G^{r}$-labeled NPSL2 (pink) at 37 °C. (C) 1D proton (aromatic) NMR spectra of $A^{2r}G^{r}$-labeled NPSL2 at different temperatures. Secondary structures were rendered using RNA2Drawer.[15]

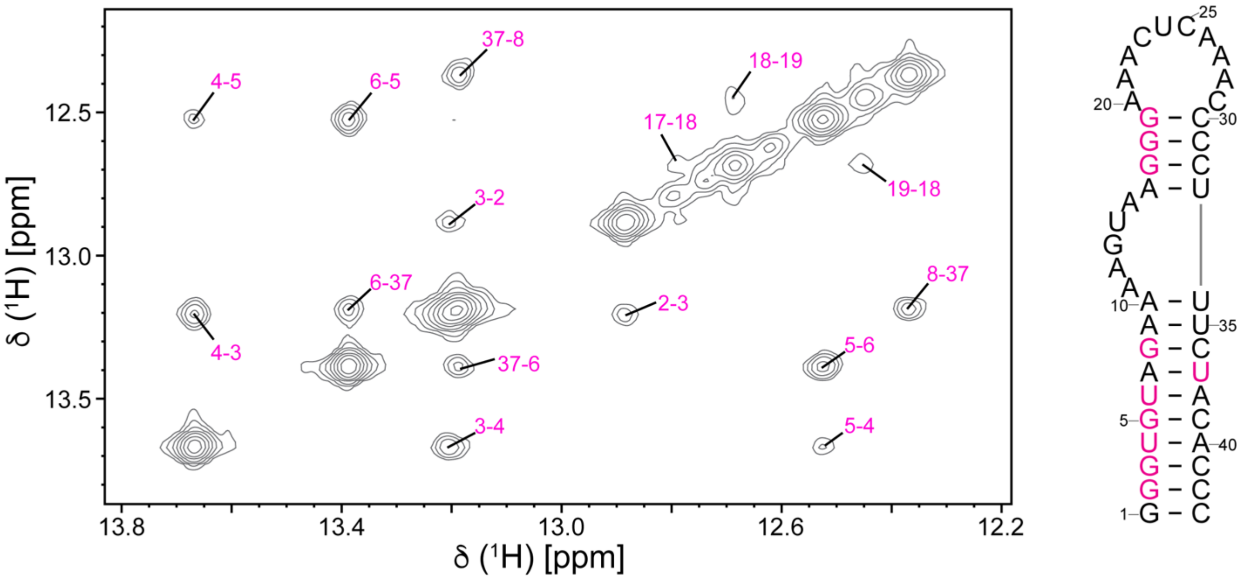


**Figure S7**

**Figure S7. Identification of base pairing in NPSL2.** Assigned imino $^{1}$H-$^{1}$H NOESY spectrum for NPSL2 (left). G's and U's assigned in the NOESY spectrum are colored magenta in the secondary structure (right). Secondary structures were rendered using RNA2Drawer.[15]

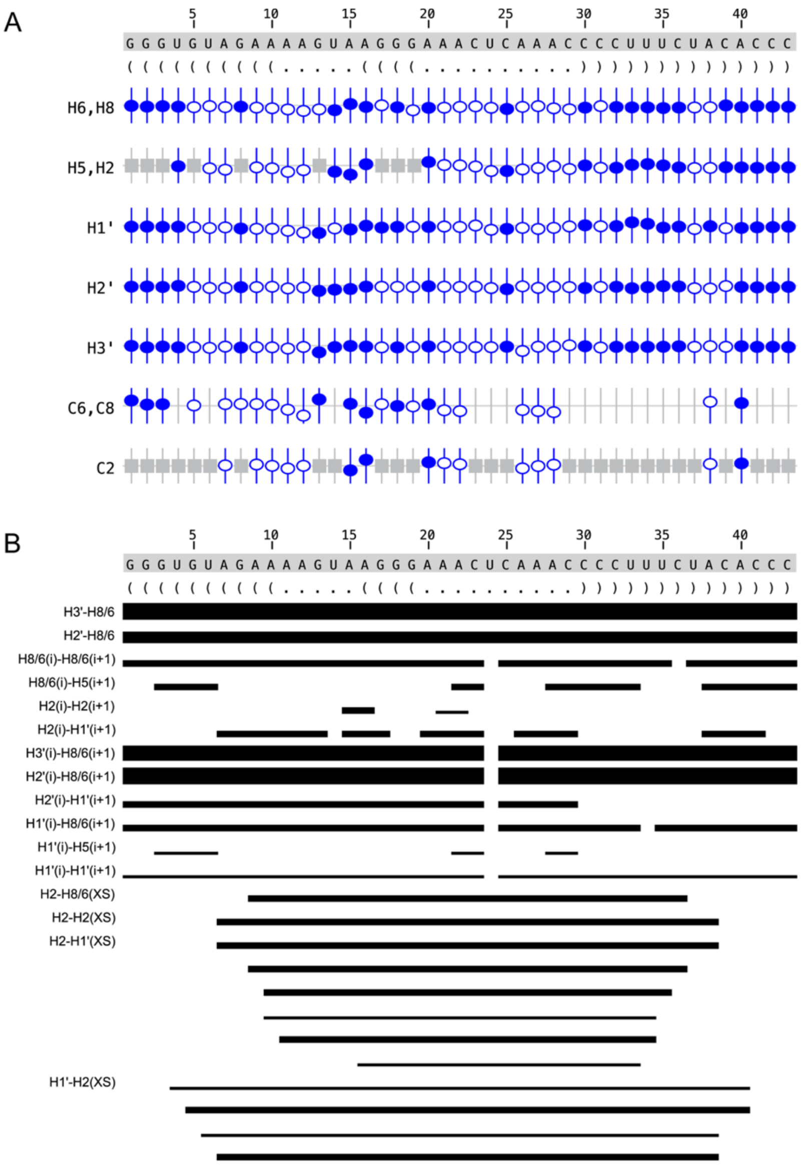


**Figure S7**

**Figure S8. Summary of NPSL2 chemical shift assignments and NOEs.** (A) Summary of the sequence, secondary structure, and assignment validation for NPSL2. The secondary structure is shown in Vienna format. Assigned chemical shifts (H6/H8, H5/H2, H1′, H2′, H3′, C6/C8, C2) are indicated with open or filled circles. The vertical offset of the circles indicates the deviation from the predicted values for that atom. Filled circles indicate that there are chemical shifts for atoms with the same set of attributes in the BMRB. Open circles indicate atoms that have a prediction, but for which no exact matches of the attributes are available in the BMRB. Grey boxes represent atoms that are not present in a given base. (B) Summary of the sequence, secondary structure, NOE connectivity, and assignment validation for NPSL2. The secondary structure is shown in Vienna format. NOE upper limit restraints for specified proton pairs used in CYANA and AMBER calculations are drawn as black bars. The thickness of the bar is representative of the strength of the measured NOE.

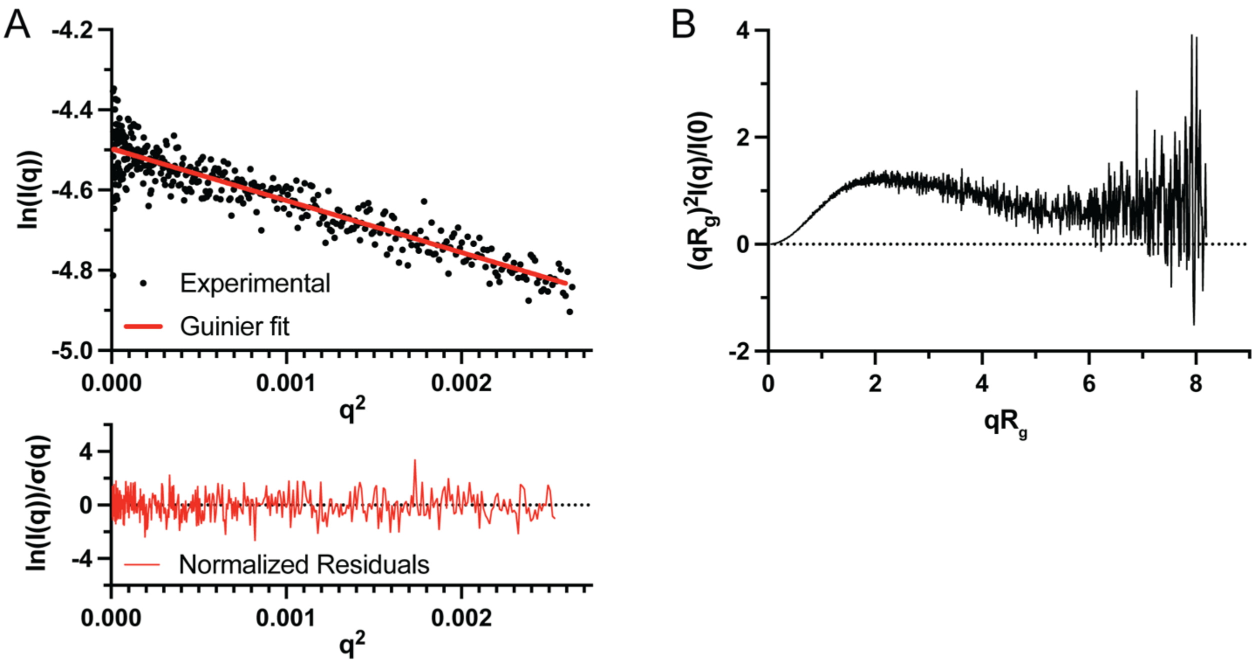


**Figure S8**

**Figure S9. Guinier analysis and Kratky plots demonstrating quality of SAXS data and suitability for further analysis.** (A) Guinier analysis of NPSL2 used to derive $R_g$ and $I(0)$ parameters (top). Normalized residuals are flat and randomly distributed about zero (bottom). (B) Dimensionless Kratky plot of NPSL2.

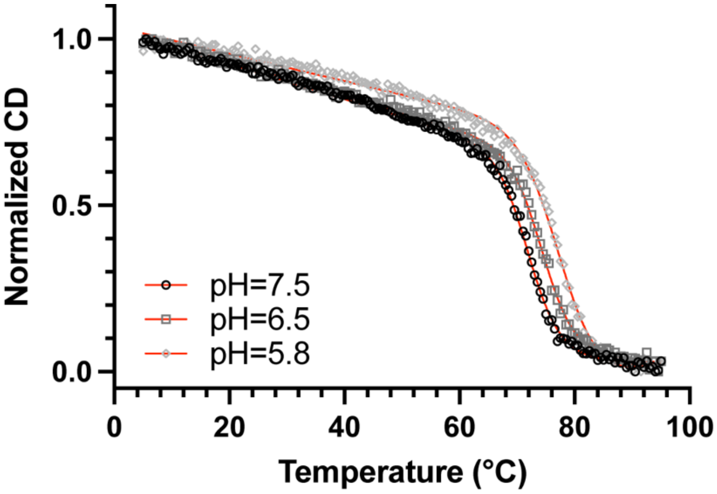


**Figure S9**

**Figure S10. NPSL2 RNA exhibits increased stability at low pH.** Thermal denaturation of NPSL2 at pH 7.5 (black circle), pH 6.5 (medium grey squares), and pH 5.8 (light grey diamonds).

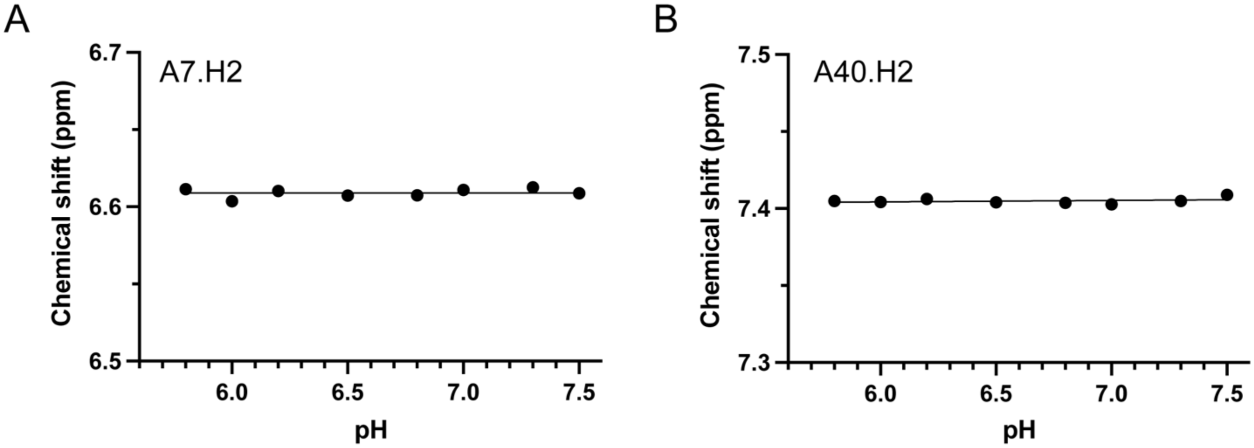


**Figure S11**

**Figure S11. A7.H2 and A40.H2 are not sensitive to pH changes.** Effect of chemical shift for (A) A7.H2 and (B) A40.H2.

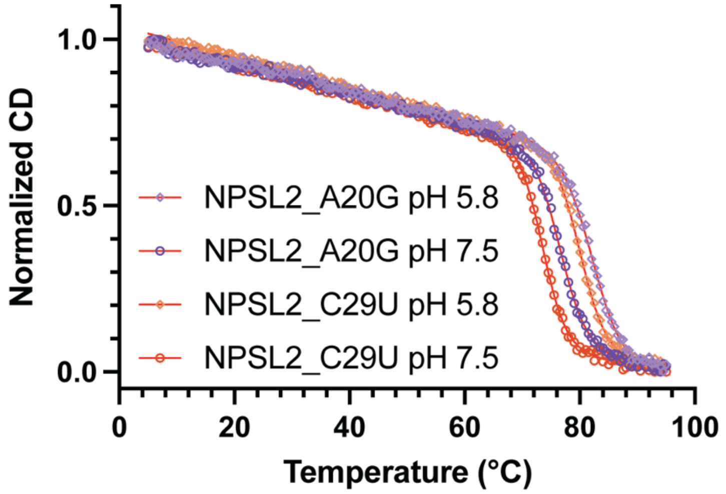


**Figure S12**

**Figure S12. Mutations that eliminate the A·C mismatch are stabilized at low pH.** CD thermal denaturation of NPSL2_A20G (purple) and NPSL2_C29U (orange) at pH 7.5 (open circles) and pH 5.8 (open diamonds).

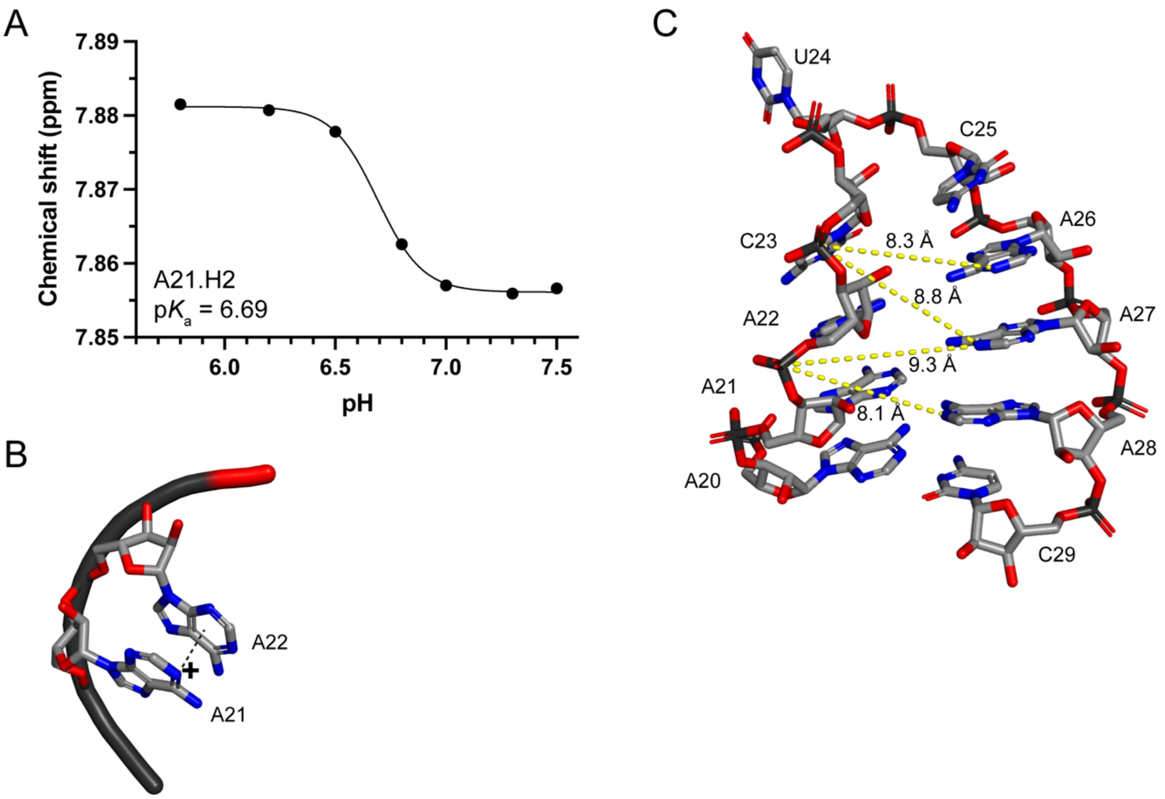


**Figure S13**

**Figure S13. Protonated adenosines in the internal loop may contribute to the stabilization of NPSL2 at low pH.** (A) Plot of the pH dependence of A21.H2 chemical shift. (B) 3D representation of cation-π interaction between protonated A21.N1 and the base of A22. (C) Favorable electrostatic interactions between phosphate groups of C23 and A22 and bases of A26, A27 and A28. The dashed yellow line indicates distance.

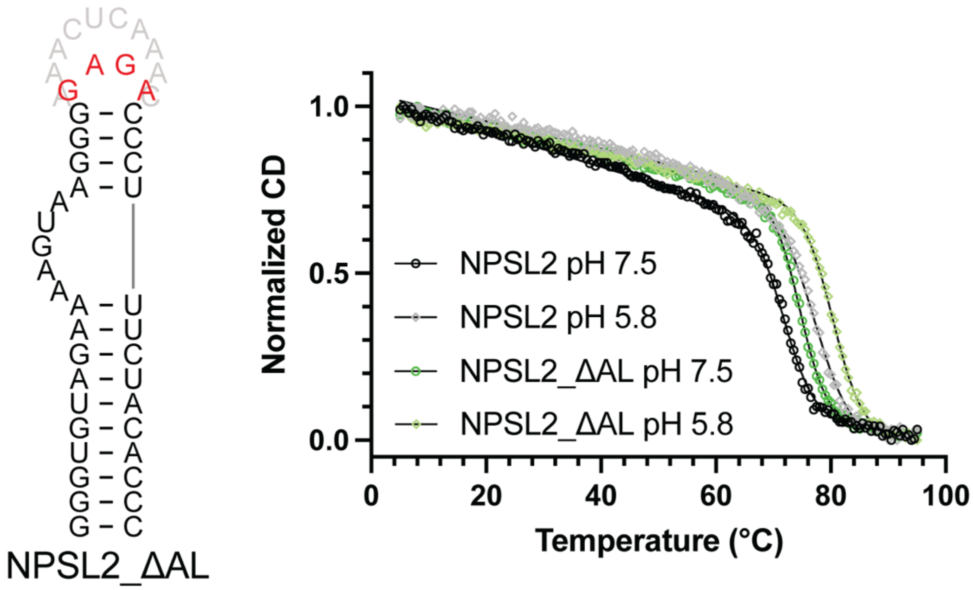


**Figure S14**

**Figure S14. Thermal stability of NPSL2 RNA lacking the apical loop increases at low pH.** Secondary structure of NPSL2_ΔAL construct (left). The apical loop residues that were removed are shaded grey and the non-native GAGA tetraloop is shown in red. Normalized thermal denaturation of NPSL2 (black) and NPSL2_ΔAL RNA (green) at pH 7.5 (open circles) and pH 5.8 (open diamonds). Secondary structures were rendered using RNA2Drawer.[15]

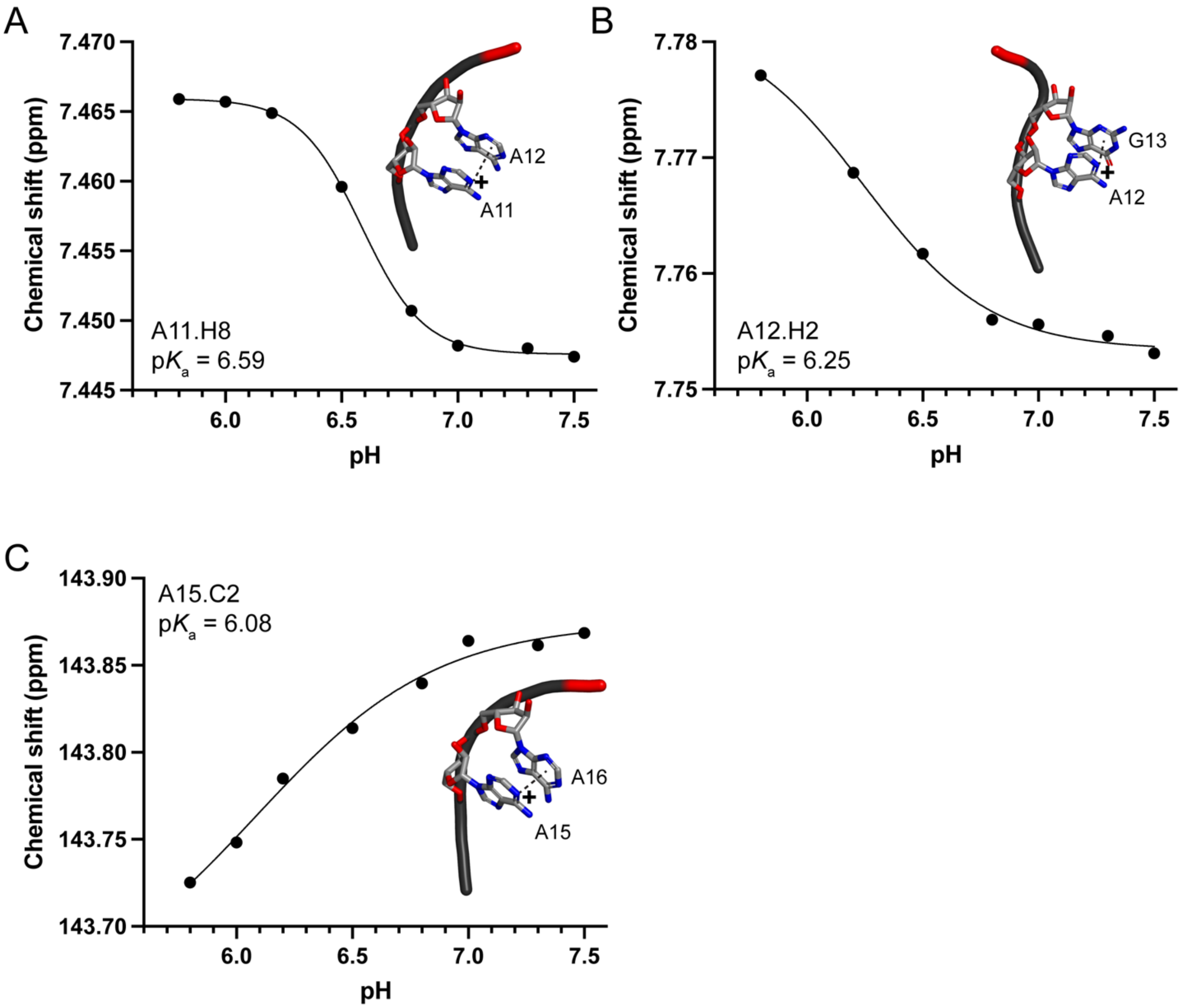


**Figure S15**

**Figure S15. Protonated adenosines in internal loop may contribute to the stabilization of NPSL2 at low pH through formation of cation-π interactions.** (A) Plot of the pH dependence of the A11.H8 chemical shift. 3D representation of a possible cation-π interaction between protonated A11.N1 and A12 (inset). (B) Plot of the pH dependence of the A12.H2 chemical shift. 3D representation of a possible cation-π interaction between protonated A12.N1 and G13 (inset). (C) Plot of the pH dependence of the A15.C2 chemical shift. 3D representation of a possible cation-π interaction between protonated A15.N1 and A16 (inset).
